# Sublethal antibiotics collapse gut bacterial populations by enhancing aggregation and expulsion

**DOI:** 10.1101/565556

**Authors:** Brandon H. Schlomann, Travis J. Wiles, Elena S. Wall, Karen Guillemin, Raghuveer Parthasarathy

## Abstract

Antibiotics induce large and highly variable changes in the intestinal microbiome even at sublethal concentrations, through mechanisms that remain elusive. Using gnotobiotic zebrafish, which allow high-resolution examination of microbial dynamics, we found that sublethal doses of the common antibiotic ciprofloxacin cause severe drops in bacterial abundance. Contrary to conventional views of antimicrobial tolerance, disruption was more pronounced for slow-growing, aggregated bacteria than for fast-growing, planktonic species. Live imaging revealed that antibiotic treatment promoted bacterial aggregation and increased susceptibility to intestinal expulsion. Intestinal mechanics therefore amplify the effects of antibiotics on resident bacteria. Microbial dynamics are captured by a biophysical model that connects antibiotic-induced collapses to gelation phase transitions in soft materials, providing a framework for predicting the impact of antibiotics on the intestinal microbiome.

## Introduction

Antibiotic drugs induce large, long-lasting, and disease-associated alterations in the composition of the intestinal microbiota [1, 2, 3]. Even at concentrations well below the minimum inhibitory levels of many bacteria, antibiotics can lead to major and highly variable changes in the gut microbiome through mechanisms that remain mysterious [2, 3, 4]. Sublethal antibiotics can also significantly alter animal physiology; the intentional growth enhancement of livestock is a well-known example that may involve microbiome-mediated pathways [2]. Low concentrations of antibiotics are often present in the environment as byproducts of unchecked agricultural and biomedical use, generating public health concerns associated with the emergence of drug resistance [5] as well as more direct impacts on human health [6]. It is therefore crucial to uncover mechanisms by which sublethal antibiotics reshape resident gut microbial communities. Understanding why particular bacterial strains are resilient or susceptible to antibiotic perturbations may allow us to predict the consequences of environmental contamination and may enable tailoring of antibiotic treatments as a therapeutic tool for manipulating the intestinal microbiome.

Conventional wisdom regarding bacterial responses to antibiotic drugs, derived largely from in vitro assays, holds that drug tolerance is facilitated by low growth rates and biofilm formation [7, 8]. Recent work suggests that microbes in the vertebrate gastrointestinal tract adopt a variety of growth and aggregation phenotypes [9, 10, 11, 12], raising the question of whether antibiotic susceptibility in the gut bears the same relationship to kinetics and physical structure as in less dynamic environments, or whether the strong mechanical activity and large fluid flows present in the intestine [13] lead to fundamentally different rules.

To investigate the in vivo response of gut bacteria to low-dose antibiotic exposure, especially the relationship between susceptibility and bacterial behavior, we conducted live imaging-based studies of larval zebrafish (Fig. 1A, 1B), spanning the entire intestinal volume with spatial and temporal resolutions not attainable in humans or other model vertebrates. We focused our study on two native zebrafish bacterial isolates, both frequently found in the intestine [14], that we identified as representing extremes of growth and aggregation phenotypes [10]. The first, *Vibrio cholerae* ZWU0020, hereafter referred to as *Vibrio*, exists in the larval zebrafish intestine primarily as dense populations of highly motile and planktonic individuals (Fig. 1C, Supplemental Movie 1). *Vibrio* grows rapidly, with an in vivo doubling time of approximately 1 hour (exponential growth rate of 0.8 *±* 0.3 1/hr) [15]. The second, *Enterobacter cloacae* ZOR0014, hereafter referred to as *Enterobacter*, primarily forms large, dense bacterial aggregates with small sub-populations of non-motile planktonic cells (Fig. 1D, Supplemental Movie 2) [16] and has an in vivo doubling time of approximately 2.5 hours (exponential growth rate of 0.27 *±* 0.05 1/hr) (Fig. S1). To delineate and quantify antibiotic responses independent of inter-bacterial competition, we studied *Vibrio* and *Enterobacter* separately in hosts that were initially raised germ-free (Materials and Methods). We assessed response dynamics of each bacterial population after treatment with the antibiotic ciprofloxacin, a broad spectrum fluoroquinolone that interferes with DNA replication by inhibiting DNA gyrase. Ciprofloxacin is widely administered therapeutically and has been used as a model antibiotic in studies of human microbiome disruption [17]. Furthermore, ciprofloxacin is often detected in environmental samples at ng/ml concentrations that are sublethal but capable of perturbing bacterial physiology [18, 19].

**Figure 1:**
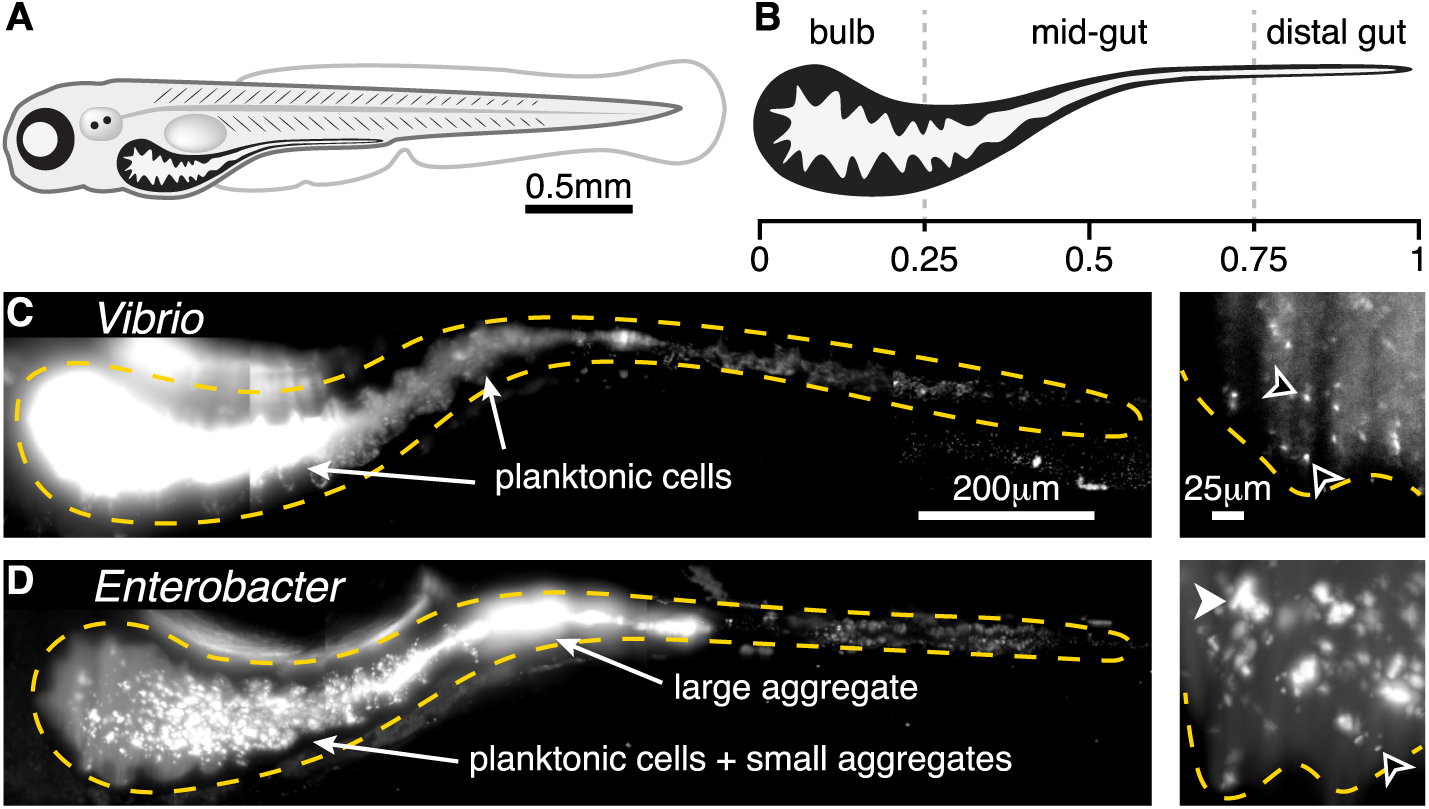
Two bacterial species show different extremes of in vivo aggregation phenotypes. A: Schematic of a zebrafish 5 days post-fertilization (dpf). B: Schematic of the larval zebrafish intestine with numbers denoting approximate fraction of gut length. C: *Vibrio cholerae* ZWU0020 in vivo. Left: a maximum intensity projection of a three-dimensional image of the full gut. Dense, bright bacteria and dimmer intestinal autofluorescence are evident. The orange dashed curve indicates a coarse outline of the gut boundary. Scale bar: 200 *µ*m. Right: a single optical plane within the anterior bulb in a fish colonized with 1:100 green fluorescent protein (GFP): dTomato (dTom)-expressing *Vibrio*, with the GFP channel shown to highlight individual microbes in the dense swarm. The orange dashed curve indicates the approximate contour of the intestinal epithelium. Black arrowheads indicate examples of single planktonic cells. Scale bar: 25 *µ*m. (See also Supplemental Movie 1) D: *Enterobacter cloacae* ZOR0014 in vivo, shown as a maximum intensity projection of the full gut (left) and a subset of the same projection in the anterior bulb (right); bacterial aggregates are evident. The black arrowhead indicates an example of a single planktonic cell; the white arrowhead indicates an example of a multicellular aggregate. Scale bars same as in (C).

As detailed below, we discovered that sublethal levels of ciprofloxacin lead to major reductions in intestinal abundance of both *Vibrio* and *Enterobacter* that could not be predicted from in vitro responses alone. In contrast to conventional wisdom, the slow-growing and highly aggregated *Enterobacter* was impacted far more severely than the fast-growing, planktonic *Vibrio*. Changes in bacterial abundances were driven primarily by clearance from the intestine by peristaltic-like fluid flow, which impacts aggregated bacteria more severely than planktonic cells. Exposure to sublethal levels of ciprofloxacin shifted both species to a more aggregated state, but for *Enterobacter* this state was unsustainable and led to population collapse and extinction. Quantitative image-derived population data motivate and are well fit by physical models originally used to describe colloidal growth and polymer gelation, implying an antibiotic-induced phase transition in bacterial community physical structure and revealing a general framework for understanding and predicting intestinal antibiotic perturbations.

## Results

### Low-dose ciprofloxacin increases bacterial aggregation and intestinal expulsion

For both *Vibrio* and *Enterobacter*, we empirically determined a ciprofloxacin dosage that induced clear changes in bacterial physiology and behavior in vitro, but that was below the apparent minimum inhibitory concentration. We first describe results of antibiotic exposure, in vitro and in vivo, for the *Vibrio* species.

From an initial survey of dose-response in rich media, we identified 10 ng/mL ciprofloxacin as an appropriate exposure for *Vibrio* populations. Growth of *Vibrio* in lysogeny broth in the presence of 1 ng/ml ciprofloxacin closely resembles that of the untreated control, while a concentration of 100 ng/ml is largely inhibitory (Fig. S2A). An intermediate concentration of 10 ng/ml leads to a stable, intermediate optical density. Viability staining (Materials and Methods) after 6 hours of incubation with 10 ng/ml ciprofloxacin identifies 30-80% of cells as alive (Fig. S3A and S3B), again consistent with this antibiotic concentration being sufficient to perturb the bacterial population without overwhelming lethality. Growth in the presence of 10 ng/ml ciprofloxacin induces marked changes in cell morphology and motility: treated cells exhibit filamentation, making them considerably longer (mean *±* std. dev. 5.3 *±* 3.1 *µ*m) than untreated *Vibrio* (2.9 *±* 0.9 *µ*m) (Fig. S2B). Swimming speed was also reduced compared to untreated cells (mean *±* std. dev. 11.4 *±* 7.2 *µ*m/s, untreated 16.9 *±* 11.1 *µ*m/s) (Fig. S2C, Supplemental Movies 3 and 4). We note also that 10 ng/ml ciprofloxacin is comparable to levels commonly measured in environmental samples [18].

While useful for illuminating the appropriate sub-lethal concentration to further examine, experiments in rich media conditions are not an optimal assay for comparison of in vitro and in vivo antibiotic treatments, as the chemical environments are likely very dissimilar. We therefore assessed effects of ciprofloxacin on bacterial populations in the aqueous environments of the flasks housing the larval zebrafish in comparison to populations in the intestines. In the flask water, as in the intestine, the only nutrients are fish-derived. Oxygen levels are comparable to those in the larval gut, due to fast diffusion and the animals’ small size. Bacteria in flask water therefore constitute a useful baseline against which to compare antibiotic impacts on intestinal populations.

*Vibrio* was associated with germ-free zebrafish at 4 days post-fertilization (dpf) by inoculation of the aqueous environment at a density of 10^6^ cells/ml (Materials and Methods) and allowed to colonize for 24 hours, which based on previous studies provides ample time for the bacterial population to reach its carrying capacity of approximately 10^5^ cells/gut [15]. Animals and their resident *Vibrio* populations were then immersed in 10 ng/ml ciprofloxacin for 24 or 48 hours, or left untreated (Fig. 2A and 2B). *Vibrio* abundances in the gut were assayed by gut dissection and plating to measure colony forming units (CFUs) (Materials and Methods). Abundances in the flask water were similarly assayed by plating. We quantified the effect of the antibiotic treatment by computing the ratio of bacterial abundances in the treated and untreated cases, resulting in a normalized abundance (Fig. 2C). After a 24 hour treatment, log_10_-transformed abundances in the flask water dropped by 0.98 *±* 0.4 (mean *±* std. dev.) compared to untreated controls, or one order of magnitude on average. In contrast, log_10_-transformed intestinal abundances showed a more severe reduction of 1.75 *±* 0.88 (Fig. 2C), or a factor of approximately 60 on average, suggesting that the intestinal environment amplifies the severity of ciprofloxacin treatment. For the 48 hour treatment, the declines in flask water and intestinal abundances were similarly severe (Fig. 2C). In terms of absolute abundances, pooled data from 24 and 48 hour treatments gives a mean *±* std. dev. of the log_10_-transformed *Vibrio* population of 3.1 *±* 0.9 (*n* = 40), compared to 4.9 *±* 0.5 (*n* = 42) for untreated specimens (Fig. 2D). Unpooled data are similar (Fig. S3E, S3F).

**Figure 2:**
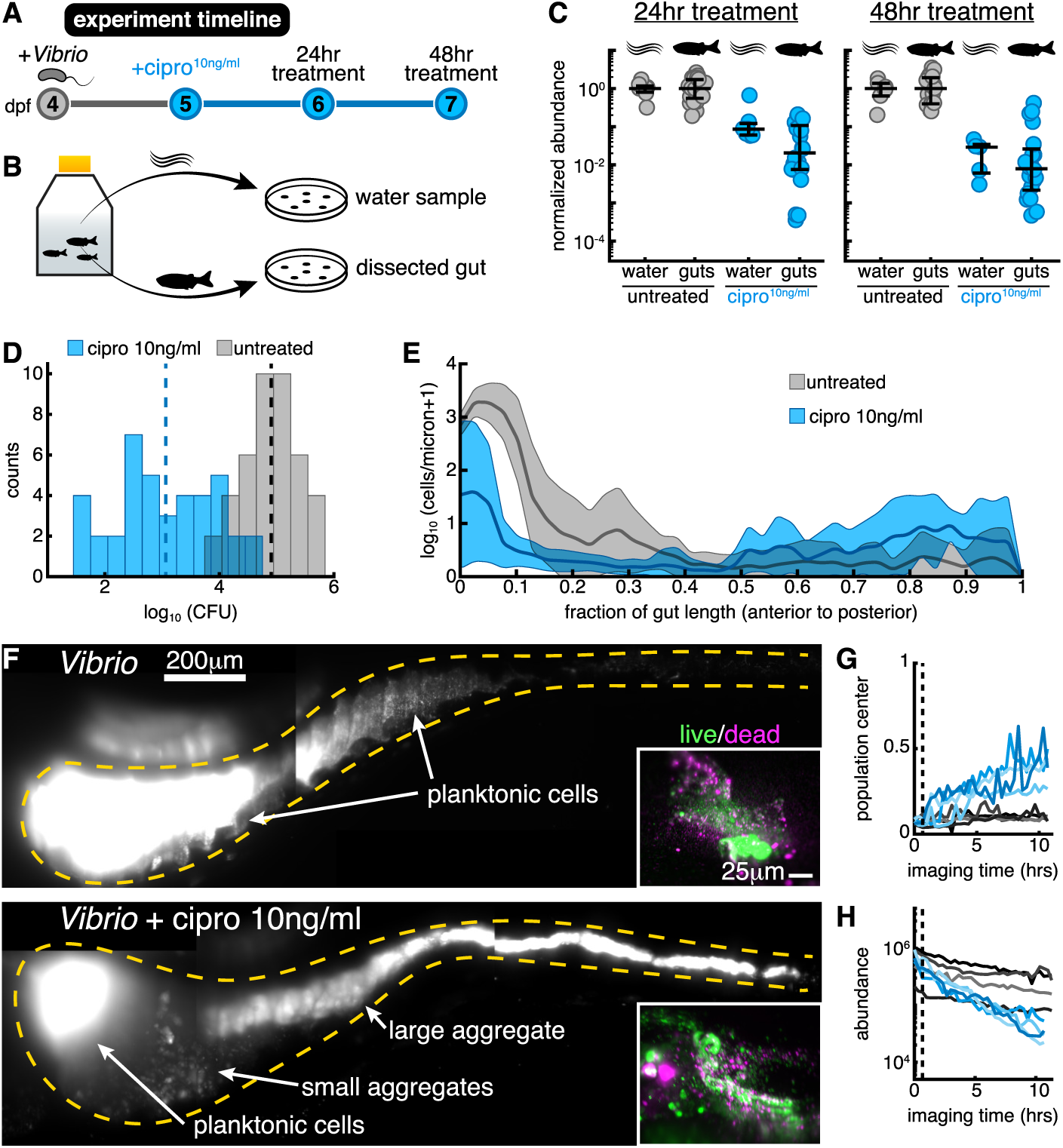
Low-dose ciprofloxacin induces *Vibrio* aggregation and expulsion in vivo. A: Schematic of the experimental timeline. B: Schematic of the sampling scheme for plating measurements. C: Normalized abundances (number of colony forming units (CFUs) scaled by untreated medians) of water and gut populations. D: Distributions of bacterial intestinal abundance of *Vibrio* mono-associated with larval zebrafish, assessed as CFUs from plating of dissected gut contents. Counts indicate the number of individual fish with a given log_10_ *Vibrio* CFUs. Dashed lines indicate the mean of each set, showing a *∼*100-fold reduction in intestinal *Vibrio* abundance in antibiotic-treated fish. E: Ensemble-averaged spatial distributions of log-transformed cell density as a function of distance along the gut axis, integrated over the perpendicular dimensions. F: Maximum intensity projections of 3D images of untreated (top) and ciprofloxacin-treated (bottom) *Vibrio* populations. Insets: Viability staining of bacteria expelled from the gut, with green and magenta indicating living and dead cells, respectively. G-H: Dynamics of in vivo *Vibrio* populations untreated (grey lines) and treated with 10 ng/ml ciprofloxacin (blue lines). G: 1D center of mass, normalized to intestine length. H: Total image-derived *Vibrio* abundance. In both (G) and (H), each curve represents a different zebrafish. Vertical dotted lines indicate the time of drug administration to the treatment cohort, *t* = 0.67 hours.

To assess the possibility that the intestine makes *Vibrio* more susceptible to ciprofloxacin-induced cell death, we embedded larval zebrafish in a 0.5% agarose gel, which allowed collection of expelled bacteria. After staining expelled bacterial cells with the viability dyes SYTO9 and propidium iodide, we imaged ejected material. We found no detectable difference between ciprofloxacin-treated and untreated populations (Fig. 2F, insets). Similarly sizeable fractions of viable and non-viable cells are evident in both ciprofloxacin-treated and untreated populations; however, co-staining of zebrafish host cells hindered exact quantification (Fig. S4). This result suggests that the ciprofloxacin-induced population decline observed in vivo occurs independent of overt cell death and is a consequence of the response of living bacteria to the intestinal environment. We further note that the dose-response of the intestinal *Vibrio* abundance (Fig. S5) mirrors the dose-response of the in vitro growth rate, implying that the larval gut does not significantly alter or concentrate ciprofloxacin. This is also consistent with the widespread use of zebrafish larvae as a pharmacological screening platform, as water soluble chemicals readily enter and leave the animal [20, 21].

To investigate the causes of ciprofloxacin’s disproportionately large impact on in vivo bacterial abundance, we used light sheet fluorescence microscopy to directly monitor *Vibrio* populations within the intestine over several hours as they responded to antibiotic exposure. Three-dimensional time-lapse imaging revealed that within hours of ciprofloxacin treatment, large numbers of bacteria became depleted from the anterior-localized planktonic and motile population (Supplemental Movies 5 and 10). Cells were instead found in the mid and distal regions of the gut, where they appeared to be condensed into large multicellular aggregates prior to being expelled from the gut altogether (Supplemental Movies 5 and 11). After 10 hours of exposure, *Vibrio* populations in ciprofloxacin-treated hosts contained large, 3D aggregates localized to the posterior of the intestine, a feature not observed in untreated controls (Fig. 2E and 2F) nor in all previous characterizations of this strain [15, 10]. We note also that in vitro, antibiotic-treated *Vibrio* does not form large aggregates (Fig. S3 and S6, Supplemental Movie 4)

**Figure 3:**
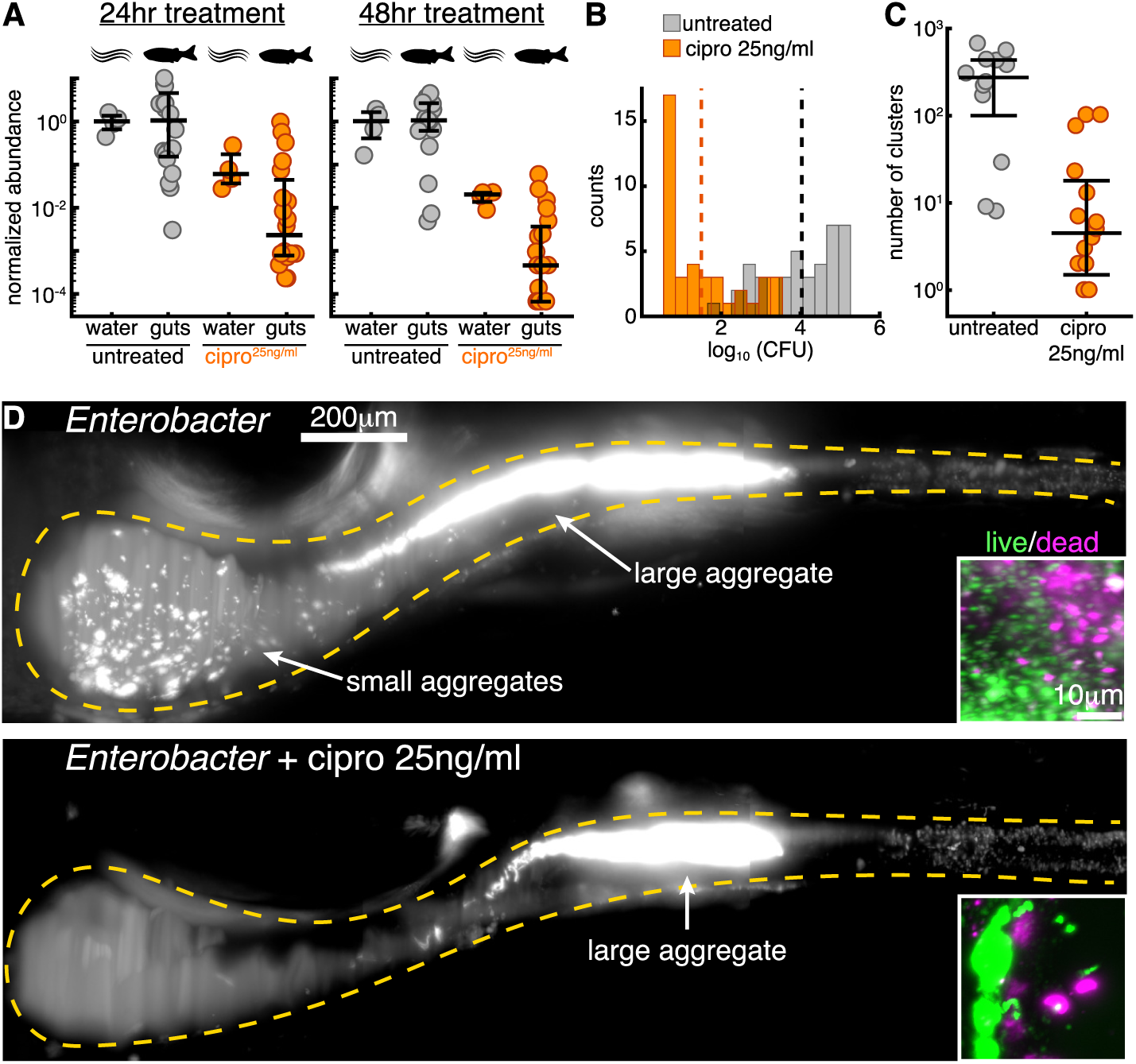
Low-dose ciprofloxacin collapses *Enterobacter* populations and suppresses small clusters in vivo. A: Normalized abundances (number of colony forming units (CFUs) scaled by untreated medians) of water and gut populations. B: Distributions of bacterial intestinal abundance of *Enterobacter* mono-associated with larval zebrafish, assessed as colony forming units (CFUs) from plating of dissected gut contents. Counts indicate the number of individual fish with a given log_10_ *Enterobacter* CFUs. Dashed lines indicate the mean of each set, showing a *∼*1000-fold reduction in intestinal *Enterobacter* abundance in antibiotic-treated fish. C: Total number of bacterial clusters in the intestine, quantified from 3D images (Materials and Methods). D: Maximum intensity projections of 3D images of untreated (top) and ciprofloxacin-treated (bottom) *Enterobacter* populations. Insets: Viability staining of bacteria expelled from the gut, with green and magenta indicating living and dead cells, respectively.

To determine whether the bacterial aggregation observed in vivo stems from a fundamentally different response to antibiotics at the single-cell level or different large-scale consequences of similar cell-level response, we generated in *Vibrio* a genetically encoded fluorescent reporter of the SOS pathway (Fig. S7, Materials and Methods), a DNA damage repair pathway induced by genotoxic agents such as ciprofloxacin [22, 23]. Genes in the SOS regulon halt replication and enable DNA repair, and also affect motility and biofilm formation [24, 19]. In vitro, we found that treatment with 10 ng/ml ciprofloxacin strongly induced *recN* -based SOS reporter activity, with a heterogeneous response across individual cells (Fig. S3C and S3D). Within the intestine, SOS reporter activity was also heterogeneous, appearing in both planktonic and aggregated cells. Planktonic cells that were SOS-positive appeared more filamented and less motile compared to SOS-negative cells within the same host (Supplemental Movie 6). The activation of the SOS reporter in vitro and in vivo by ciprofloxacin (Supplemental Movie 6 and Fig S3C and S3D) suggests that in both cases a canonical SOS response is involved in the perturbation of *Vibrio* physiology.

Together, these data begin to reveal a mechanism by which the intestine amplifies the effect of low-dose ciprofloxacin. Individual *Vibrio* cells first undergo an SOS response that is associated with changes in cellular morphology and behavior. In the context of the mechanical activity of the intestine, these molecular and cellular-level changes then give rise to population-level aggregation and spatial reorganization throughout the entire length of the intestine, with the population shifting its center of mass posteriorly (Fig. 2G, *n* = 4 per case). This process culminates in the expulsion of large bacterial aggregates from the host, causing a precipitous decline in total bacterial abundance (Fig. 2H).

### Low-dose ciprofloxacin suppresses small cluster reservoirs associated with intestinal persistence

In contrast to *Vibrio*, *Enterobacter* is slower growing, non-motile, and naturally forms dense aggregates within the zebrafish intestine. *Enterobacter* populations have an in vivo growth rate of 0.27 *±* 0.05 h^−1^ (mean *±* std. dev, Fig. S1), compared to 0.8 *±* 0.3 h^−1^ for *Vibrio* [15]. Based on conventional notions of antibiotic tolerance, we hypothesized that *Enterobacter* would be less affected by ciprofloxacin treatment than the fast growing, planktonic *Vibrio*. However, as detailed below, we found this prediction to be incorrect; *Enterobacter* exhibits an even greater response to low-dose ciprofloxacin.

We first established in vitro that 25 ng/ml ciprofloxacin produces similar effects on *Enterobacter* growth as did 10 ng/ml exposure on *Vibrio*. With the identical inoculation procedure used for *Vibrio*, log_10_-transformed *Enterobacter* abundance in the flask water dropped by 1.2 *±* 0.4 (mean *±* std. dev.) compared to untreated controls after 24 hours, and dropped by 1.8 *±* 0.2 after 48 hours (Fig. 3A). These values match well the values for *Vibrio*: 0.98 *±* 0.37 for 24 hours, 1.81 *±* 0.5 for 48 hours. Assays in rich media show a similarly reduced density between the two species (Fig. S8) and an even lesser degree of cell death and damage in vitro for *Enterobacter* as compared to *Vibrio*, with a viable fraction of approximately 95% (Fig. S9A and S9B). As with *Vibrio*, in vitro growth measurements and viability staining both imply that low-dose ciprofloxacin treatment of *Enterobacter* induces growth arrest rather than widespread lethality.

Strikingly, low-dose ciprofloxacin treatment of fish colonized with *Enterobacter* (Materials and Methods) resulted in even greater reductions in abundance than in the case of *Vibrio*, with the majority of populations becoming nearly or completely extinct during the assay period (Fig. 3A and 3B). Inoculation, treatment, dissection, and plating were performed as for *Vibrio* (Materials and Methods). Compared to untreated controls, log_10_-transformed intestinal abundances were reduced by 2.3 *±* 1.1 after 24 hours, and by 3.2 *±* 1.0 after 48 hours (Fig. 3A). These reductions in intestinal abundances greatly exceeded the reductions of bacterial abundances in the flask water (Fig 3A). In terms of absolute abundances, pooled data from 24 and 48 hour treatments gives a mean *±* std. dev. of the log_10_-transformed *Enterobacter* population of 1.5 *±* 1.0 (*n* = 40), compared to 4.0 *±* 1.0 (*n* = 39) for untreated specimens (Fig. 3B); unpooled data are similar (Fig. S9C and S9D).

Live imaging of intestinal populations at single time points revealed approximately 40% of treated hosts to be devoid or nearly devoid of *Enterobacter*, consistent with the plating-based measurements. In hosts that contained appreciable bacterial populations we observed a clear difference between treated and untreated specimens: *Enterobacter* populations in ciprofloxacin-treated hosts contained fewer small bacterial clusters and fewer individual planktonic cells than untreated controls (Fig. 3C and 3D). We quantified this distinction using computational image analysis to identify each cluster (Materials and Methods), defining a single cell as a cluster of size one. Bacterial populations in ciprofloxacin-treated animals contained *∼*80x fewer clusters than untreated animals (Fig. 3C). Viability staining showed that there were no obvious differences in the viable fractions of bacteria expelled from the intestines of untreated and treated hosts (Fig. 3D, insets, Fig. S10). As with *Vibrio*, these observations suggested that the reduction in *Enterobacter* ’s intestinal abundance was independent of cell death.

Previous studies of other naturally aggregated bacterial species have revealed that large bacterial aggregates are highly susceptible to expulsion from the gut [15, 25]. To establish whether this is also the case for *Enterobacter* in the absence of low-dose ciprofloxacin treatment, we performed time-lapse 3D imaging (Materials and Methods). Indeed, in 2 out of 5 hosts imaged for 3.5 hours each, we observed events in which the largest bacterial aggregate was abruptly expelled from the intestine (Fig. 4A and Supplemental Movie 7). These timelapse movies also showed clear examples of cluster aggregation (Supplemental Movie 8), in which single cells and small aggregates appear to come together and fuse, a process that is likely due to the rhythmic intestinal contractions that occur between frames. Importantly, smaller aggregates and planktonic cells that preferentially localize to the intestinal bulb are relatively undisturbed during these expulsion events, save for a few clusters that become incorporated into the large mass during its transit (Supplemental Movie 7).

**Figure 4:**
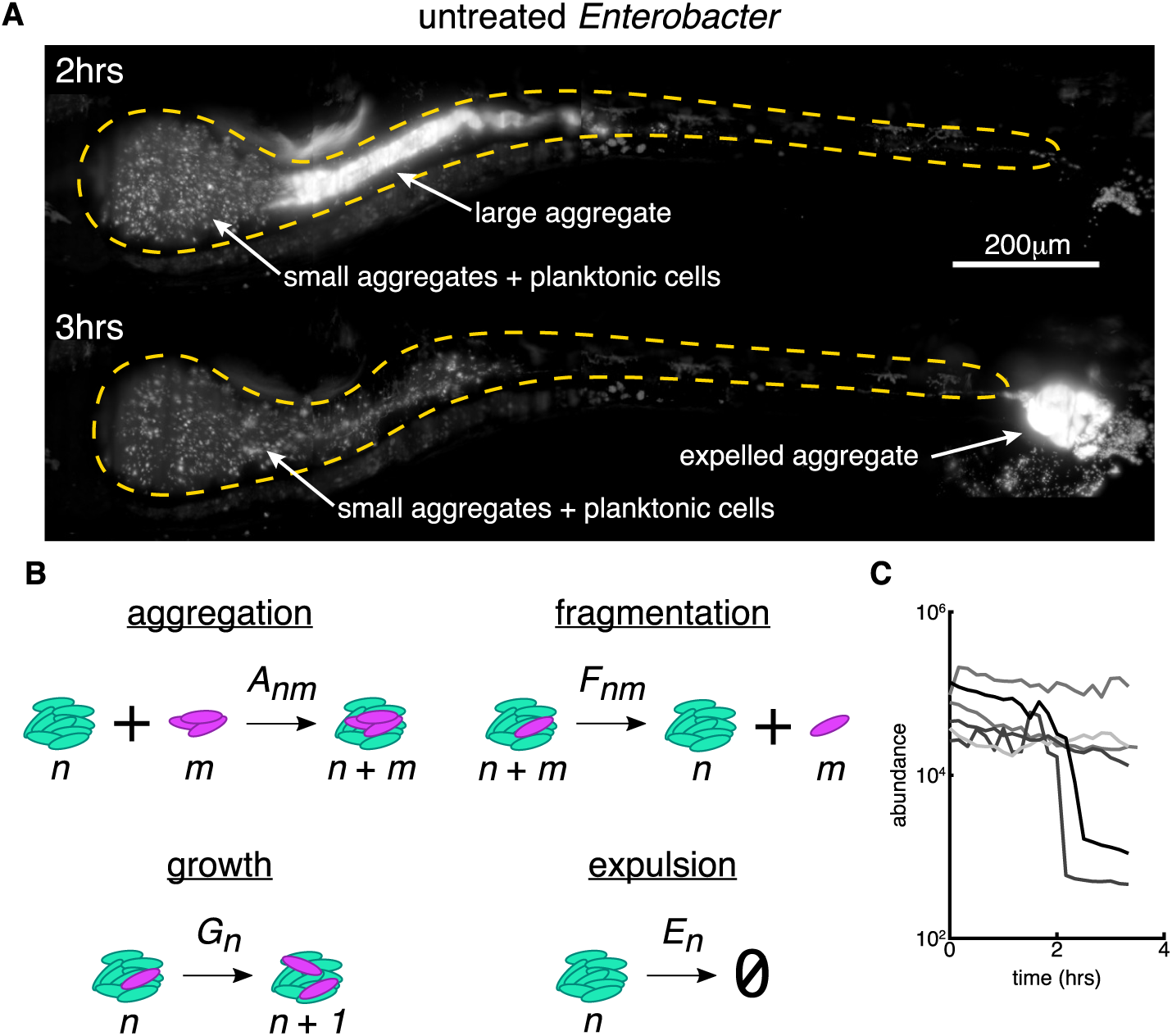
Small bacterial clusters are required for recovery after large expulsion events. A: Maximum intensity projections of untreated *Enterobacter* populations before (top, *t* = 2 hours from the start of imaging) and after (bottom, *t* = 3 hours) an expulsion event (See also Supplemental Movie 5). Scale bar = 200 *µ*m. B: Schematic of a kinetic model of bacterial cluster dynamics, illustrating its four constituent processes. C: Image-derived time-series of *Enterobacter* abundance in five untreated hosts showing sporadic large expulsion events.

Our observations suggest an explanation of how low-dose ciprofloxacin can lead to dramatic drops in *Enterobacter* abundance that moreover illuminates the more general question of how naturally aggregating bacterial species can persist in the vertebrate gut in spite of transport-driven expulsion. We provide both a qualitative and a quantitative description of the relevant dynamics, beginning with the following conceptual model: single cells of *Enterobacter* replicate to form small clusters, which then aggregate to form larger clusters under the influence of intestinal flow. Large clusters are transported by the rhythmic contractions of the gut [15, 25, 26] and are stochastically expelled from the host [15, 25]. The individual bacteria and small clusters that remain within the intestine serve as a reservoir that reseeds the next population, and the process of replication, aggregation, and expulsion repeats. Therefore, persistence within the intestine requires processes that generate single cells or small clusters, otherwise transport will eventually lead to extinction. This reseeding could take the form of (i) immigration of new cells from the environment, (ii) passive fragmentation of clusters, or (iii) active fragmentation in which single cells break away from a cluster surface during cell division. Immigration from the environment likely occurs even in established populations, but measurements in larval zebrafish suggest very low rates of immigration [27]. We therefore suspected that more robust mechanisms must promote persistence. Supporting the active fragmentation mechanism, we found in untreated hosts examples of *Enterobacter* populations that contain an abundance of single cells, a single large aggregate, and a lack of mid-sized aggregates (Fig. S9E). Following low-dose ciprofloxacin treatment, the planktonic cell reservoir associated with resilience to intestinal transport is depleted (Fig. 3C), most likely due to stalled *Enterobacter* division (Fig. S8), leading to collapse of the resident bacterial population (Fig. 3A and 3B).

### A quantitative model of bacterial cluster dynamics

To solidify and test our conceptual picture, we developed a predictive mathematical model of bacterial cluster dynamics. We describe the framework of the model, its validation, and general insights it provides into perturbations and population stability. Drawing on ideas from non-equilibrium statistical mechanics and soft matter physics, we constructed a general kinetic model that describes the time evolution of a collection of bacterial clusters with varying sizes, illustrated schematically in Fig. 4B. We posit that four processes govern cluster dynamics: aggregation, fragmentation, growth, and expulsion. Each is described by a kernel that specifies its rate and size dependence: (1) aggregation of a cluster of size *n* and a cluster of size *m* occurs with rate *A_nm_*; (2) fragmentation of a cluster of size *n* + *m* into clusters of size *n* and *m* occurs with rate *F_nm_*; (3) growth (due to cell division) of a cluster of size *n* occurs with rate *G_n_*; (4) expulsion (removal by intestinal transport) of a cluster of size *n* occurs with rate *E_n_*. Note that condensation of the population into a single massive cluster poises the system for extinction, for any nonzero *E_n_*. The model is non-spatial and is inspired by well established frameworks for nucleation and growth phenomena such as polymer gelation and colloidal aggregation [28]. For example, sol-gel models describe a transition between dispersed individual units (“sol”) and a system-spanning connected network (“gel”) in materials capable of polymerization. In the thermodynamic limit of infinite system size, the model can be studied using standard analytic techniques [28]. However, unlike polymer solutions and other bulk systems for which the possible number of clusters is effectively unbounded, our intestinal populations are constrained to have at most a few hundred clusters (Fig. 3C), necessitating the use of stochastic simulations (Materials and Methods).

In its general form, the model encompasses a wide range of behaviors that can be encoded in the various functional forms possible for the rate kernels *A_nm_ F_nm_*, *G_n_*, and *E_n_*. Based on our observations and theoretical considerations elaborated in the Materials and Methods section, we made the following assumptions: (1) the rate of aggregation between two clusters is independent of their size, *A_nm_* = *α*; (2) fragmentation occurs only by separation of single cells and with a rate that is independent of cluster size, *F_nm_* = *β* for *m* = 1 and *F_nm_* = 0 otherwise; (3) growth is logistic with a global carrying capacity, *G_n_* = *rn*(1 *− N/K*) with *N* the total number of cells, *r* the per capita growth rate, and *K*, the carrying capacity; (4) expulsion is independent of cluster size, *E_n_* = *λ*. This model contains as special cases various simple models of linear polymers [29] and also resembles recent work modelling chains of *Salmonella typhimurium* cells in the mouse gut [30]. As discussed in the Materials and Methods section, these choices constitute the minimal model consistent with theoretical constraints and experimental phenomenology. More complex models are of course possible, but the requisite increase in the number of adjustable parameters would result in a trivial but meaningless ability to fit the observed data.

Even with the assumptions described above, the model needs 5 parameters: rates of aggregation, fragmentation, growth, and dispersal, and a global carrying capacity. However, all of these parameters can be set by experimentally derived values unrelated to cluster size distributions. We measured *Enterobacter* ’s per capita growth rate by performing time-lapse imaging of initially germ-free hosts that had been exposed to *Enterobacter* for only 8 hours, capturing the exponential increase of a small founding population (Fig. S1, Supplemental Movie 9), yielding *r* = 0.27 *±* 0.05 hr^−1^ (mean *±* std. dev across *n* = 3 hosts). We identified expulsion events as abrupt collapses in *Enterobacter* abundance from time-lapse images (Fig. 3C, Supplemental Movie 7) and set the expulsion rate equal to the measured collapse rate, *λ* = 0.11 *±* 0.08 hr^−1^ (mean *±* standard error, assuming an underlying Poisson process (Materials and Methods)). The model can be simulated to provide the mean and variance of the log_10_-transformed abundance distribution at a given time for a given set of parameters. Using this approach, we fit static bacterial abundance measurements from dissection and plating at 72 hours post-inoculation (Materials and Methods) to determine the carrying capacity, *K*, and the ratio of fragmentation and aggregation rates, *β/α*. As discussed in the Materials and Methods section, the cluster dynamics should depend primarily on the ratio of *β/α* rather than either rate separately. This yielded log_10_ *K* = 5.0 *±* 0.5 and log_10_ *β/α* = 2.5 *±* 0.4.

The model therefore allows a parameter-free prediction of the size distribution of *Enterobacter* aggregates, plotted in Fig. 5A together with the measured distribution derived from three-dimensional images, averaged across 12 untreated hosts. The two are in remarkable agreement. We also plot, equivalently, the cumulative distribution function *P* (size *> n*), the probability that a cluster will contain greater than *n* cells, again illustrating the close correspondence between the data and the prediction and validating the model. We emphasize that no information about the cluster size distribution was used to estimate any of the model parameters. We further note that the cluster size distribution is a stringent test of the model’s validity. Other cluster models predict different forms, typically with steep tails [29, 30]. The linear chain model of [30], for example, leads to an exponential distribution of cluster sizes that does not match the shallower, roughly power-law form of our data.

**Figure 5:**
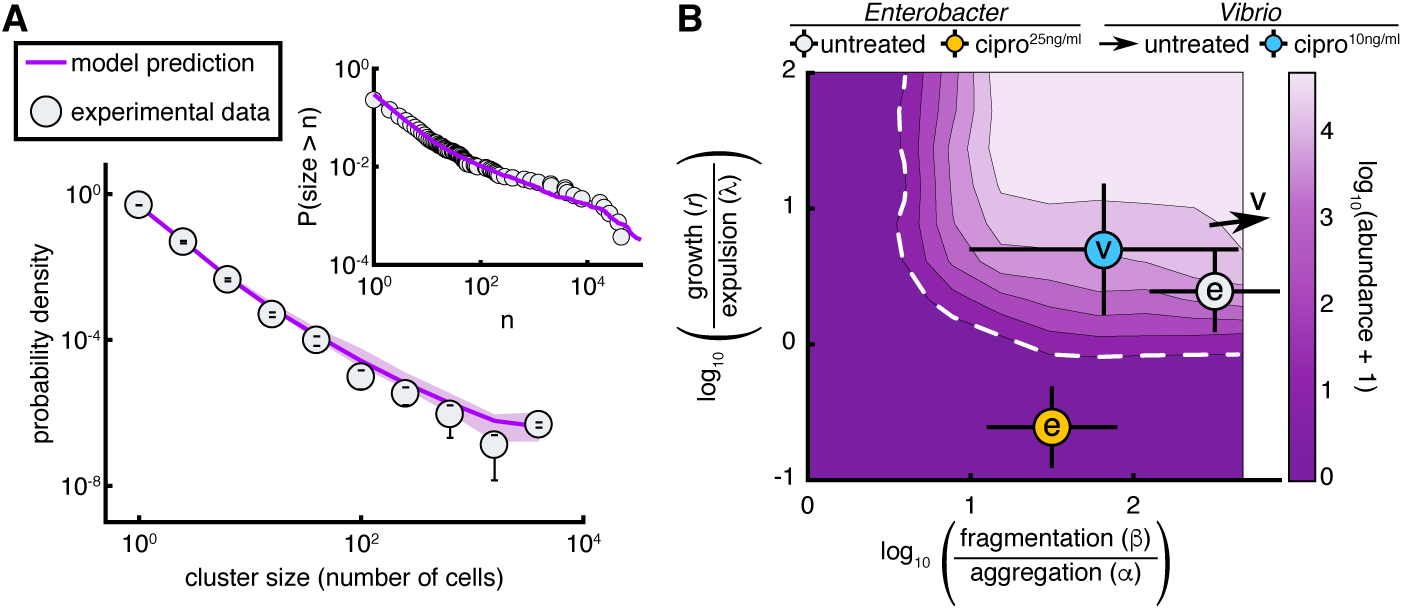
A stochastic kinetic model predicts bacterial cluster sizes and generates a phase diagram for in vivo abundance. A: The distribution of image-derived *Enterobacter* cluster sizes (grey circles) along with the prediction of our stochastic model (purple line). There are no free parameters in the fit; values were fixed by abundance, growth, and expulsion rate measurements independent of cluster size. Parameters: *r* = 0.27 hr^−1^, *λ* = 0.11 hr^−1^, *α* = 0.1 hr^−1^, *β* = 10^1.5^ hr^−1^, *K* = 10^5^. Error bars on experimental data are standard deviations across hosts. Shaded confidence intervals for the model prediction are bounds from parameter uncertainties. Inset: The same experimental data and model plotted without binning as a reverse cumulative distribution. B: Phase diagram of the log-transformed abundance, (log_10_(*N* + 1)), showing the extinction transition (white dashed line). From best fit parameter estimates, the in vivo state of untreated *Enterobacter* is overlaid as a grey circle, and 25 ng/ml ciprofloxacin-treated *Enterobacter* as an orange circle; both circles are marked with “e”. Untreated *Vibrio* is located off the scale, indicated by the arrow, 10 ng/ml ciprofloxacin treated *Vibrio* is overlaid as a cyan circle marked with “v”. The doses for *Enterobacter* and *Vibrio* were established to be approximately equivalent in vitro. Parameters: *λ* = 0.11 hr^−1^, *α* = 0.1 hr^−1^, *K* = 10^5^ were fixed; *r* and *β* were varied on logarithmic grids.

### The abundance phase diagram and extinction transition

Our kinetic model provides insights into the consequences of low-dose antibiotic perturbations on gut bacterial populations. We consider a general phase diagram of possible growth, fragmentation, aggregation, and expulsion rates, and then situate *Enterobacter* in this space. For simplicity of illustration, we consider a two-dimensional representation with one axis being the ratio of the fragmentation and aggregation rates, *β/α*, and the other being the ratio of the growth and expulsion rates, *r/λ* (Fig. 5B). As noted above and in the Materials and Methods section, the model in the regime studied should depend on the ratio *β/α* rather than on *β* or *α* independently. However, the roles of *r* and *λ* are not simply captured by their ratio. The expulsion rate nonetheless provides a scale to which to compare the growth rate, *r*, and we plot Fig. 5B using *r/λ* calculated for fixed *λ* = 0.11 hr^−1^, the measured value. For completeness, we show a three-dimensional *r, λ, β/α* phase diagram as Figure S11E and S11F. We numerically calculated the steady state phase diagram of the model (Materials and Methods) and show in Figure 5B the mean log-transformed abundance, 〈log_10_(*N* + 1)〉. The regime of extinction (*N* = 0) is evident (dark purple, with dashed white boundary).

The data-derived parameter values place untreated intestinal *Enterobacter* fairly close to the extinction transition (Fig. 5B). An antibiotic-induced growth rate reduction of approximately 5x is sufficient to cross the boundary to the *N* = 0 regime (i.e. to extinction), moving downward in Fig. 5B. This growth rate reduction, or an equivalent increase in death rate, reflects the conventional view of antibiotic effects. An approximately 300x reduction in the balance between fragmentation and aggregation spurs an alternative path to extinction, moving leftward in Fig. 5B, reflecting a distinct mechanism resulting solely from changes in physical structure. The extinction transition in this direction corresponds to the condensation of the population into a single cluster, reminiscent of gelation phase transitions in polymer systems. As described above, low-dose ciprofloxacin causes a strong reduction in the number of small bacterial clusters, lowering *β* and possibly also *r* if fragmentation and individual cell division are linked. Conservatively assuming an equal effect along both axes, and fitting simulations to the 24 hour treatment abundances (Materials and Methods), we find that the antibiotic reduces *r* and *β/α* by *∼*10x. This drives the bacterial system through the phase boundary and well into the extinction regime (Fig. 5B, orange circle), consistent with our observations.

In contrast to *Enterobacter*, treatment of *Vibrio* with ciprofloxacin does not lead to widespread extinction after 48 hours, suggesting that treated populations either lie safely at a new steady state away from the extinction boundary, or are close enough to the transition so that dynamics are slow. To estimate model parameters for ciprofloxacin-treated *Vibrio*, we performed a two parameter fit of (*β/α, r*) to the 24 hour treatment abundances. Because of *Vibrio*’s large population size (*∼* 10^5^ clusters), we modified the stochastic simulation procedure using a tau-leaping algorithm (Materials and Methods, Fig. S12). We indeed find ciprofloxacin-treated *Vibrio* is located close to but safely inside the extinction boundary (Fig. 5B). Untreated *Vibrio* populations show no appreciable multicellular aggregation and are located off-scale far to the upper-right side of the phase diagram (Fig. 5B, arrow).

## Discussion

We have discovered that sublethal levels of a commonly used antibiotic can reduce the intestinal abundance of bacterial populations much more severely than would be predicted from in vitro responses, and that this amplification is a consequence of drug-induced changes to the bacterial groups’ spatial architecture. Contrary to conventional notions of antibiotic tolerance, largely derived from in vitro studies, reductions in bacterial abundances were greater for the slow-growing, aggregated *Enterobacter* species than for the fast-growing, planktonic *Vibrio*. Live imaging revealed drug-induced increases in bacterial cohesion that, coupled to gut mechanical activity, lead to the expulsion of viable bacterial cells. The microscopic details of this cohesion, likely involving cell wall characteristics, mechanical compression by the gut wall and fluid flows, and perhaps intestinal mucus rheology, remain to be explored.

Notably, the underlying processes of bacterial aggregation and host intestinal transport are found throughout the animal kingdom, suggesting a general relevance beyond zebrafish that may explain, for example, data on weak antibiotics having strong effects on mammalian microbiomes [2, 3]. Of course, chemical perturbations in more anatomically complex animals or non-gnotobiotic animals that house hundreds of resident bacterial species will undoubtedly involve additional processes beyond those uncovered here. We note, however, that responses to intestinal flow will influence bacterial population dynamics regardless of ecological complexity, and that our choice of model bacterial species spans the extremes of highly planktonic and highly cohesive strains, further implying generality. In the larval zebrafish, enhanced bacterial susceptibility to transport leads to expulsion from the gut. In larger or more complex intestines this may take the form of displacement from one region to a more distal region, with a corresponding shift in local nutrients or competitors, in addition to expulsion from the gut altogether.

The concentrations of ciprofloxacin examined here are commonly found in environmental samples, indicating a potentially widespread perturbation of animal gut microbiota due to antibiotic contaminants. In addition, the expulsion of live, antibiotic-exposed bacteria from animal intestines through the aggregation-based processes described here suggests a potential mechanism for enhanced spread of antibiotic resistance. This possibility is bolstered by our observation that in addition to aggregation, ciprofloxacin-treated cells undergo an active SOS response, which has been shown to promote mutation and horizontal gene transfer [31, 32, 33]. Together, these observations underscore recent concerns about the public health risk posed by antibiotic contaminants in the environment [6].

Our biophysical model of aggregation, fragmentation, growth, and expulsion describes our data well and provides testable predictions. It is remarkable, given the chemical and morphological complexity of even the larval zebrafish gut, that such a minimal model can accurately predict emergent properties such as the size distribution of bacterial aggregates. That this works is an indication of the power of theories of soft condensed matter physics, whose generality may prove useful in understanding the gut microbiome. Furthermore, our model supplies a framework for a quantitative understanding of gut microbial homeostasis in general. Like recent work modelling antibody-mediated enchaining of *Salmonella* cells in the mouse gut [30], our model implies that the physical processes of bacterial cluster formation and fragmentation play central roles in large-scale microbiota stability. We suggest that our cluster-dynamics model, validated by quantitative agreement between predictions and in vivo data (Fig. 5A), may prove useful in less tractable host species such as mice and humans. Without live imaging or non-invasive sampling, it is challenging to estimate kinetic properties of microbial populations, such as aggregation rates. However, advances in histological sample preparation [34] can preserve bacterial aggregates and yield cluster size distributions; inverting our model, such distributions can reveal the underlying in vivo bacterial kinetics.

Regarding antibiotics, the main prediction of our model is that naturally aggregated, slow growing bacteria will be impacted more severely than fast growing, planktonic species by equivalent low-dose antibiotic perturbations. This is contrary to conventional wisdom that links bacterial tolerance to reduced growth and increased aggregation [7, 8], which stems from studies of antibiotic exposure in static or well-mixed environments. We find that in the intestine, where bacteria can be removed through fluid flow, there exist critical values of aggregation, fragmentation, growth, and expulsion rates, beyond which sustainable colonization becomes impossible (Fig. 5B). Naturally aggregated and slow-growing species are situated closer to this extinction phase boundary and are therefore more easily driven to population collapse by low-dose antibiotic perturbations. Intriguingly, new meta-omics methods [9] can be used to estimate in vivo growth rates of mammalian gut microbes, which would be interesting to correlate with antibiotic responses. We note in addition that inter-bacterial competition in the gut can be influenced by clustering and susceptibility to intestinal transport [15, 25], suggesting that competition outcomes could be altered by antibiotic treatment if changes in aggregation properties are different for different species. A final prediction of our model is that intestinal transport, which has been linked to microbiota composition [13], will influence the effects of low-dose antibiotic perturbations on microbial community composition. Combining pharmacological manipulations of intestinal transport with antibiotic treatments may therefore lead to novel strategies for precision engineering of the gut microbiome.

## Acknowledgements

We thank Rose Sockol and the University of Oregon Zebrafish Facility staff for fish husbandry. Research was supported by an award from the Kavli Microbiome Ideas Challenge, a project led by the American Society for Microbiology in partnership with the American Chemical Society and the American Physical Society and supported by The Kavli Foundation. Work was also supported by the National Science Foundation under Awards 1427957 (R.P.) and 0922951 (R.P.), the M.J. Murdock Charitable Trust, and the National Institutes of Health (http://www.nih.gov/), under Awards P50GM09891 and P01GM125576-01 to K.G. and R.P., F32AI112094 to T.J.W., and T32GM007759 to B.H.S. The content is solely the responsibility of the authors and does not represent the official views of the NSF, National Institutes of Health, or other funding agencies. This work benefited from access to the University of Oregon high performance computer, Talapas.

## Materials and Methods

### Animal care

All experiments with zebrafish were done in accordance with protocols approved by the University of Oregon Institutional Animal Care and Use Committee and following standard protocols [35].

### Gnotobiology

Wild-type (AB*×*TU strain) zebrafish were derived germfree (GF) and colonized with bacterial strains as previously described [36] with slight modifications. Briefly, fertilized eggs from adult mating pairs were harvested and incubated in sterile embryo media (EM) containing ampicillin (100 *µ*g/ml), gentamicin (10 *µ*g/ml), amphotericin B (250 ng/ml), tetracycline (1 *µ*g/ml), and chloramphenicol (1 *µ*g/ml) for 6 hours. Embryos were then washed in EM containing 0.1% polyvinylpyrrolidone-iodine followed by EM containing 0.003% sodium hypochlorite. Sterilized embryos were distributed into T25 tissue culture flasks containing 15 ml sterile EM at a density of one embryo per milliliter and incubated at 28 to 30*^◦^*C prior to bacterial colonization. Embryos were sustained on yolk-derived nutrients and were not fed during experiments. For bacterial mono-association, bacteria were first grown overnight in lysogeny broth (LB) with shaking at 30*^◦^*C and were prepared for inoculation by pelleting 1 ml of culture for 2 min at 7,000*×g* and washing once in sterile EM. Bacterial strains were individually added to the water column of single flasks containing 4-day-old larval zebrafish at a final density of 10^6^ bacteria/ml. For antibiotic treatment, drugs were added at the indicated working concentration directly to flask containing animals that had been colonized 24 hours prior.

### Bacterial strains and culture

*Vibrio cholerae* ZWU0020 and *Enterobacter cloacae* ZOR0014 were originally isolated from the zebrafish intestine [14]. Fluorescently marked derivatives of each strain were previously generated by Tn*7* -mediated insertion of a single constitutively expressed gene encoding dTomato [16]. We note that all plating- and imaging-based experiments performed in this study were done using fluorescently marked strains, which carry a gentamicin resistance cassette, with the exception of experiments in which fluorescent dyes were used to assess viability of cells. Archived stocks of bacteria were maintained in 25% glycerol at −80*^◦^*C. Prior to experiments, bacteria were directly inoculated from frozen stocks into 5 ml LB media (10 g/L NaCl, 5 g/L yeast extract, 12 g/L tryptone, 1 g/L glucose) and grown for *∼*16 hours (overnight) shaking at 30*^◦^*C.

### Generation of a fluorescent SOS reporter

To identify a suitable promoter within the *Vibrio* ZWU0020 genome (https://img.jgi.doe.gov/m/, IMG genome ID: 2522572152) for creation of a genetically encoded fluorescent DNA-damage ‘SOS’ reporter, we scanned the upstream regions of each gene for consensus gammaproteobacterial ‘SOS boxes’ (CTGTN_8_ACAG) that serve as binding sites for the repressor LexA (Fig. S7A and S7B) [22]. Of the genes identified, the promoter of the gene *recN* (IMG gene ID: 2705597027) was an ideal candidate for three main reasons: 1) it contains multiple SOS boxes (2 consensus and 2 with 2 mismatches), which is an arrangement that is potentially associated with tight/graded regulation [23]; 2) the *recN* promoter is highly conserved among closely related *V. cholerae* strains as well as other non-*Vibrio* gammaproteobacterial lineages, suggesting that *recN* is a bona fide representative of the SOS response; and 3) *recN* is one of the most highly expressed genes in response to DNA damaging agents in both *E. coli* and *V. cholerae* [37, 38], likely due to its multiple near-consensus −10 promoter sequences.

We rationally designed a *recN* -based fluorescent SOS reporter by fusing the 100bp *recN* promoter region to an open reading frame (ORF) encoding superfolder green fluorescent protein (sfGFP) (Fig. S7C). In addition, we incorporated an epsilon enhancer and consensus Shine-Dalgarno sequence within the 5’ untranslated region (UTR) to help ensure robust translation of the reporter gene [16, 39, 40], and incorporated the synthetic transcriptional terminator L3S2P21 into the 3’ UTR [41]. We built the construct using polymerase chain reaction (PCR) and synthetic oligonucleotides. Primer WP97 (containing the *recN* promoter and 5’ UTR; 5’-TGAATGCATTAAAAGTGACCAAAAAATTTTACCTGAGTGACTTTACTGTATAAAGAAACAGTATAAACTGTTTAAACATACAGTATTGGTTAATCATACAGGTGCAAACTTAACTTTATCAAGGAGACTAAATCATGAGCAAGGGCGAGGAGCT-3’) and primer WP98 (containing the 3’ UTR; 5’-TGAACTAGTAAAACGAAAAAAGGCCCCCCTTTCGGGAGGCCTCTTTTCTGGAATTTTTATCACTTGTACAGCTCGTCCATG-3’) were used to PCR-amplify sfGFP from the source plasmid pXS-sfGFP [16]. Engineered restriction sites flanking the amplicon (NsiI and SpeI) were then used to insert the construct into a variant of the Tn*7* delivery vector pTn*7* xKS, which also harbors a constitutively expressed *dTomato* gene for tracking all bacterial cells (Fig. S7D) [16]. The resulting dual-reporter construct was then inserted into the ZWU0020 genome as previously described [16]. To verify reporter activity, disk diffusion assays were performed on agar plates with the genotoxic agent mitomycin C and, as a control, the cell wall-targeting beta-lactam antibiotic ampicillin (Fig. S7E). Mitomycin C induced robust expression of sfGFP whereas ampicillin did not.

### In vitro characterization of antibiotics

#### Growth kinetics

Growth kinetics of bacterial strains in vitro were measured using a FLU-Ostar Omega microplate reader. Prior to growth measurements, bacteria were grown overnight in 5 ml LB media at 30*^◦^*C with shaking. The next day, cultures were diluted 1:100 into fresh LB media with or without the indicated antibiotic and dispensed in quadruplicate (200 *µ*l/well) into a sterile 96-well clear flat-bottom tissue culture-treated microplate. Absorbance at 600 nm was then recorded every 30 min for *∼*16 hours at 30*^◦^*C with shaking. Growth rates were estimated by fitting a logistic growth curve to OD values, starting at manually defined points marking the end of lag phase.

#### Viability

Cultures of *Vibrio* ZWU0020 or *Enterobacter* ZOR0014 were grown overnight in LB at 30*^◦^*C with shaking. The next day, 1:100 dilutions were made in fresh LB media containing either ciprofloxacin (*Vibrio*: 10 ng/ml, *Enterobacter* : 25 ng/ml) or no drug. Cultures were incubated at 30*^◦^*C with shaking for 6 hours prior to being stained using a LIVE/DEAD BacLight Bacterial Viability Kit according to manufacturer’s instructions. Culture/stain mixtures were diluted 1:10 in 0.7% saline and imaged using a Leica MZ10 F fluorescence stereomicroscope equipped with a 2.0X objective and a Leica DFC365 FX camera. Images were captured using standard Leica Application Suite software. Bacteria were identified in images with intensity-based region finding following difference of gaussians filtering. Cells stained in both SYTO9 and propidium iodide were identified as overlapping regions in the two color channels. Analysis code was written in MATLAB.

#### Cell length and swimming speed

Dense overnight cultures of *Vibrio* ZWU0020 were diluted 1:100 in fresh LB media alone or with 10 ng/ml ciprofloxacin and incubated at 30*^◦^*C with shaking for 4 h. Bacteria were then imaged on a Nikon TE2000 inverted fluorescence microscope between a slide and a coverslip using a 60X oil immersion objective and a Hamamatsu ORCA CCD camera (Hamamatsu City, Japan). Movies were taken within 60 seconds of mounting at an exposure time of 30 ms, resulting in a frame rate of 15 frames/sec, and had a duration of approximately 7 seconds. Bacteria in the resulting movies were identified with intensity-based region finding and tracked using nearest-neighbor linking. Analysis code was written in MATLAB. Five movies were taken per treatment case. For untreated length analysis, *n* = 2291 bacteria were quantified; for ciprofloxacin-treated length analysis, *n* = 963. For untreated speed analysis, *n* = 833 bacteria; for ciprofloxacin-treated speed analysis, *n* = 531.

#### Vibrio SOS reporter activity

*Vibrio* ZWU0020 carrying the fluorescent SOS reporter was grown overnight in LB at 30*^◦^*C with shaking. The next day, 1:100 dilutions were made in fresh LB media containing either 10 ng/ml ciprofloxacin, 400 ng/ml mitomycin C, 10 *µ*g/ml ampicillin, or no drug. Cultures were then grown overnight (*∼*16 h) at 30*^◦^*C with shaking. The next day, cultures were diluted 1:43 in 80% glycerol (as an immobilizing agent) and imaged with a Nikon Eclipse Ti inverted microscope equipped with an Andor iXon3 888 camera using a 40x objective and 1.5x zoom. Bacteria were identified in images with gradientbased region finding, using a Sobel filter, following difference of gaussians filtering. Analysis code was written in MATLAB. As expected, the two DNA targeting drugs, ciprofloxcain and mitomycin C, induced the SOS response in subpopulations of cells, while the cell-wall targeting drug ampicillin did not. In computing SOS-positive fractions, filamented cells were counted as single cells.

### Culture-based quantification of bacterial populations

Dissection of larval guts was done as described previously [42]. Dissected guts were harvested and placed in a 1.6 ml tube containing 500 *µ*l sterile 0.7% saline and *∼*100 *µ*l 0.5 mm zirconium oxide beads. Guts were then homogenized using a bullet blender tissue homogenizer for *∼*25 seconds on power 4. Lysates were serially plated on tryptic soy agar (TSA) and incubated overnight at 30*^◦^*C prior to enumeration of CFU and determination of bacterial load. Typically an overnight incubation is sufficient to recover all viable cells; however, we note that ciprofloxacin treatment results in delayed colony growth on agar plates (likely due to growth arrest induced by DNA-damage). We empirically determined that, in the case of ciprofloxacin treatment, an incubation period 72 hours was required for complete resuscitation of viable cells on agar plates. For all culture-based quantification of bacterial populations in this study, the estimated limit of detection is 5 bacteria/gut and the limit of quantification is 100 bacteria/gut. Plating data plotted are pooled from a minimum of two independent experiments. Samples with zero countable colonies on the lowest dilution were set to the limit of detection prior to plotting and statistical analysis. Enumeration of flask water abundances by plating was performed identically to gut abundances, including the 72 hour incubation period.

#### Comparing antibiotic treatments between intestinal populations and flask water populations

To compare the effect of ciprofloxacin on populations in the intestine and in the flask water, we normalized treated abundances by the corresponding untreated median abundance (Fig. 2C and 3A). To control for variation in untreated bacterial dynamics between weekly batches of fish, we performed the normalization within each batch. Unnormalized data is available in the Supplemental Data File.

### Light sheet fluorescence microscopy of live larval zebrafish

#### Imaging intestinal bacteria

Live imaging of larval zebrafish was performed using a custom-built light sheet fluorescence microscope previously described in detail [43]. Larvae are anesthetized with MS-222 (Tricane) and mounted into small glass capillaries containing 0.5% agarose gel by means of a metal plunger. Larvae are then suspended vertically in an imaging chamber filled with embryo media and anesthetic and extruded out of the capillary such that the set agar plug sits in front of the imaging objective. The full intestine volume (*∼*1200 *×* 300 *×* 150 microns) is imaged in four subregions that are registered in software after imaging. The imaging of a full intestine volume sampled at 1-micron steps between *z*-planes is imaged in *∼*45 seconds. Excitation lasers at 488 and 561 nm wavelengths were tuned to a power of 5 mW prior to entering the imaging chamber. A 30 ms exposure time was used for all 3D scans and 2D movies. Time lapse imaging was performed overnight, except for the 3.5 hour imaging of *Enterobacter* (Fig. 3C), which occurred during the day.

#### Viability staining of expelled aggregates

Germ-free larval zebrafish were colonized with wild type *Vibrio* or *Enterobacter* (without fluorescent markers) for 24 hours and then mounted into agarose plugs using small glass capillaries identically to the imaging procedure (above). Individual capillaries were suspended into isolated wells of a 24-well tissue culture plate filled with embryo media containing anesthetic or anesthetic + ciprofloxacin (10 ng/ml for *Vibrio*, 25 ng/ml for *Enterobacter*) and the larvae were extruded from the capillaries. Fish remained mounted for 24 hours, during which expelled bacteria remained caught in the agarose plug. After treatment, fish were pulled back into the capillaries and transferred to smaller wells of a 96 well plate containing embryo media, anesthetic, and the LIVE/DEAD BacLight Bacterial Viability stains SYTO9 and propridium iodide. Fish were stained according to kit instructions, with the exception of the incubation period being extended from 15 to 30 min to account for potential issues with the aggregate nature of the cells [44]. Following staining, fish were pulled again into the capillaries and transferred to the light sheet microscope for imaging. As shown in Figures S4 and S10, zebrafish cells stain in addition to bacterial cells, precluding accurate quantification of viable fractions.

### Image analysis

Bacteria were identified in three-dimensional light sheet fluorescence microscopy images using a custom MATLAB analysis pipeline previously described [43, 10], with minor changes. In brief, small objects (single cells and small aggregates) are identified using difference of Gaussians filtering. False positives are rejected with a combination of intensity thresholding (mostly noise) and manual removal (mostly host cells). Large aggregates are identified with a graph cut algorithm [45] that is seeded with either an intensity-based mask or a gradientbased mask. The average intensity of a single cell is estimated as the mean intensity of small objects, which is then used to estimate the number of cells contained in larger clusters by normalizing the total fluorescence intensity of each cluster. Spatial distributions along the length of the gut are computed using a manually drawn line drawn that defines the gut’s center axis.

### Kinetic model and stochastic simulations

#### Choosing rate kernels

Our approach to choosing the size dependence of the rate parameters was to pick the simplest kernels consistent with key experimental observations. The first key observation, made in past work [43, 15], was that in between the expulsion of large aggregates population growth is well-described by a deterministic logistic function. Therefore, we chose a logistic growth kernel. The second key observation was that we occasionally encountered populations consisting of just a single, large aggregate and many single cells (Fig. S9E), which suggests that active fragmentation of single cells, most likely during growth phases, is the dominant fragmentation process. This notion is supported by time-lapse images of initial growth (Supplemental Movie 9) that depicts the creation of single cells during growth, in addition to the growth of three dimensional aggregates. Based on these observations, we made the assumption that single cell fragmentation is the sole fragmentation process, leading to what is known in other contexts as a “chipping” kernel [28]. Beyond the chipping assumption, we had little evidence that informed how single cell fragmentation depends on the size of the aggregate, so we opted for the simplest choice of a constant, size-independent rate. Similarly for aggregation and expulsion, the size dependence of the rates is likely determined by complicated and uncharacterized fluid mechanical interactions of bacterial clusters in peristaltic-like flow, which we parsimoniously replace with a simple constant kernel for both processes. In aggregated populations, since it is only the loss of the largest clusters (of size *O*(*K*)) that significantly impacts the system, we expect that it is the expulsion rate for these largest clusters that matters, rather than how the expulsion rate scales with cluster size. To test this notion, we ran simulations in which the expulsion rate scaled as a power of the cluster size, *n*, according to *λ*(*n*) = *λ*(*n/K*)*^ν^*, and varied the exponent *ν*. This ansatz keeps the expulsion rate of clusters of size *K* fixed for all values of *ν*. The result is that the cluster size distribution does not change within uncertainty values (Fig. S13), indicating that this approximation is valid.

For reference, we note that with these choices the model can be summarized by the following Smoluchowski equation, which describes the time evolution of the concentration of clusters of size *n*, *c_n_*, in the thermodynamic limit of infinite system size:

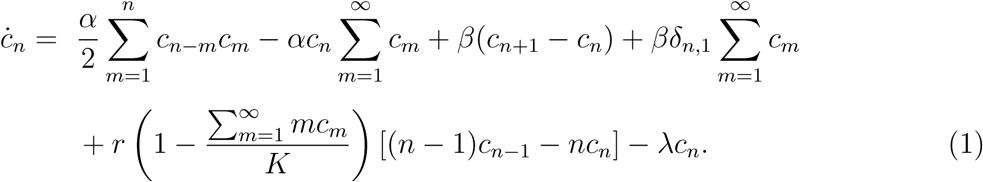

The four rate parameters are *α* (aggregation), *β* (fragmentation), *r* (growth), and *λ* (expulsion), and *K* is the carrying capacity. In the last term of the first line, *δ_n,_*_1_ is the Kronecker delta with second argument equal to 1. Of note, the first line of equation (1), containing just aggregation and fragmentation terms, was previously studied as a model of polymer chains and was shown to exhibit interesting non-equilibrium steady states and scaling behaviors that are due to the breaking of detailed balance by the chipping kernel [29]. In our system detailed balance is also broken, but for a different reason: our “monomers”—single cells—are alive and self-replicating.

#### Simulations

As each zebrafish intestine contains at most a few hundred bacterial clusters, finite size effects and stochasticity impact cluster statistics, so we implemented the model as a hybrid deterministic-stochastic simulation that follows the time evolution of individual clusters. Gillespie’s direct method [46] was used to simulate stochastic aggregation, fragmentation, and expulsion events. Growth was treated as deterministic. Once the time until next stochastic reaction, *τ*, was determined according to the Gillespie algorithm, integration was performed with the Euler method from time *t* to *t* + *τ* using a time step Δ*t* = min(*τ,* 0.1 hr).

To simulate *Vibrio* populations, direct stochastic simulation becomes intractable due to the large number of clusters (*∼* 10^5^ single cells). We therefore implemented a modified tau-leaping algorithm [47] that facilitates large simulations. We opted for a straightforward fixed *τ* method and empirically determined that a value of *τ* = 0.001 h produced no observable differences in cluster size and abundance distributions compared to direct stochastic simulation (Supp Fix X A,B).

All simulations were written in MATLAB and code is available at https://github.com/ bschloma/gac.

### Parameter inference

The kinetic model presented in the main text has 5 parameters: rates of growth, expulsion, aggregation, and fragmentation, along with an overall carrying capacity. As discussed in the main text, we directly measured *Enterobacter* ’s growth rate and expulsion rate through timelapse imaging. The uncertainty of the expulsion rate was estimated by the standard error, using the previously validated assumption that the expulsion of large aggregates follows a Poisson process [15]:

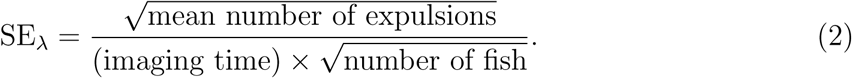

For the remaining parameters, we developed a method to infer them from the distribution of abundances obtained from dissection and plating assays. In a regime where aggregation and fragmentation are fast compared to expulsion, we expect the system to locally reach a quasi-steady state in between expulsions of the largest aggregates. As such, we expect cluster statistics to depend primarily on the ratio of fragmentation to aggregation, *β/α*, rather than on each rate independently. This confirmed in simulations (Fig. S11A and S11B). Therefore, the number of parameters to be estimated is reduced to two: *β/α* and *K*.

#### Untreated Enterobacter

We fixed *α* = 0.1 hr^−1^ and performed a grid search in *β* and *K* on a logarithmic grid, simulating the model multiple trials for each pair of (*β, K*). The number of trials decreased with increasing *β*, from 1000 to 10. Each simulation started from 10 single cells and ran for a simulated time of 64 hours, modeling our 72 hour colonization data with an 8 hour colonization window. To model static host-host variation, we drew each carrying capacity from a log-normal distribution with a standard deviation of 0.5 decades. This is the standard deviation of the untreated *Vibrio* abundance distribution (Fig. S3F), which is an appropriate measure of static host-host variation because untreated *Vibrio* does not form large aggregates and therefore does not experience large, stochastic population collapses due to aggregate expulsion. We then compared the mean (*µ*) and variance (*σ*) of the simulated, log-transformed abundances log_10_(*N* + 1) with the values for our plating data (*µ̂* and *σ̂* respectively), quantifying error using

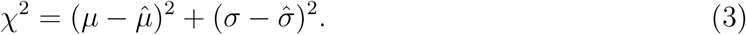

A heat map of *χ*^2^ shows well-defined edges for the minimum values of the fit parameters (Fig. S11C). However, the inference is poorly constrained for carrying capacities larger than 10^5^ and for log_10_ *β/α* greater than 2.5. This poor constraint is due primarily to the insensitivity of the abundance distribution to increasing values of these parameters. For example, moving to the far right side of the abundance phase diagram in Fig. 5B, the contours become flat in *β/α*.

To further constrain our estimates, we place upper bounds on these parameters with simple estimates of physical limits. To bound the carrying capacity, we note that a larval zebrafish intestine have a volume of roughly 1 nl, or 10^6^ *µ*m^3^. Taking the volume of a bacterium to be roughly 1 *µ*m^3^, we estimate a maximum bacterial load of 10^6^ cells, consistent with the largest *Vibrio* abundances (Fig. S3F). As we find no *Enterobacter* populations above 10^5.5^, and in our simulations we draw carrying capacities from a log-normal distribution with a standard deviation of half a decade, we constrained our best fit estimate to log_10_ *K* = 5.0. To bound the fragmentation rate, *β*, we considered the time-lapse movie that showcases the greatest degree of cluster fragmentation observed (Supplemental Movie 9). This movie depicts the initial growth phase, in which both the size of aggregates and the number of single cells increase. Because the aggregates visibly grow in size, we know that the fragmentation rate must be bounded by the absolute growth rate of the population, *β < rN*; if the fragmentation rate were larger, cells would break off of the aggregate faster than they would be produced by cell division, and the aggregates would shrink in size. Taking, roughly, *r ∼* 10^−1^ and *N ∼* 10^3^ (Fig. 4D), we estimate that *β <* 10^2^, or, with *α* = 10^−1^, *β/α <* 10^3^. With this bound, we constrain our best fit estimate to log_10_ *β/α* = 2.5. We took the uncertainties of the best fit estimates, *σ*_log10_ *_K_* and *σ*_log10_ *_β/α_*, to be the inverse of the local curvatures of *χ* at the best fit values: 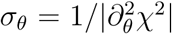, for *θ* = log*_θ_*_10_*K*, log_10_ *β/α*, resulting in *σ*_log10_ *_K_* = 0.5 and *σ*_log10_ *_K_* = 0.4.

#### Ciprofloxacin-treated Enterobacter

To estimate the change in *Enterobacter* ’s parameters upon antibiotic treatment, we conservatively assumed equal effects on growth and fragmentation/aggregation and modeled treatment parameters as *r′* = *∊r* and *β′* = *∊β*. We then performed a single parameter grid search of *∊* values, ranging from 10^−1.75^to 10^−0.5^. We modeled the antibiotic treatment as a parameter quench with a 6 hour buffer time, in which the antibiotics entered the intestine and began to take action on the bacteria. The value of 6 hours was chosen based on the *Vibrio* time series data. Each simulation was initialized with a cluster configuration drawn randomly from the imaging-derived dataset of actual untreated *Enterobacter* populations. The parameters *r*, *λ*, and *K* were set to their best fit or measured values, *α* was again fixed at 0.1 hr^−1^, and *r* and *β* were both scaled by the same factors of *∊*. We then ran simulations for a modified simulation time 24 *−* 6 = 18 hours and fit the mean and standard deviation of shifted log-transformed abundances measured in the 24 hour treatment plating assays. A plot of *χ*^2^ vs *∊* shows a clear minimum at *∊* = 10^−1^ (Fig. S11D).

#### Untreated Vibrio

Untreated *Vibrio* populations are comprised of almost entirely single cells and therefore represent an extreme limit of the kinetic model. In this regime, fragmentation is so thorough that even dividing cells immediately separate and there is no appreciable aggregation. Because multicellular clusters are extremely rare, our data are insufficient to extract numerical estimates of model parameters. However, one can estimate a lower bound for the fragmentation rate, *β*, by equating it to the total growth rate, *rN*, where *N* is the total population size; i.e. clusters do not grow without fragmenting. This estimate yields *β* 2: 10^5^. For the expulsion rate, if we assume the same rate as *Enterobacter* (positing unchanged intestinal mechanics), we obtain *r/λ ∼* 7 These values place untreated *Vibrio* off-scale in the phase diagram of Fig. 5B.

#### Ciprofloxacin-treated Vibrio

We performed a two-parameter fit to (*β/α*, *r*), using the measured expulsion rate for *Enterobacter* (*λ* = 0.11 h^−1^ and the typical untreated *Vibrio* abundance for a carrying capacity of *K ∼* 10^5^. We observed that in approaching the extinction transition from above, simulated abundance distributions transition from unimodal to bimodal in shape, with a peak emerging near *N* = 0 representing populations that suffered large, abrupt collapses. As such, fitting just the mean and variance as was done for *Enter-obacter* produced inaccurate estimates. Therefore, we implemented full maximum likelihood estimation using 100 simulated replicates to estimate the likelihood. While the fit to treated *Vibrio* resulted in less-constrained parameter estimates in the *r−β* plane compared to the Enterobacter fit, it did yield a clear maximum (Fig. S12C) and a best-fit abundance distribution that matched experimental data within uncertainties (Fig. S12D). Like with *Enterobacter*, we can attempt to assess the validity of this model by comparing the now-parameter-free prediction of the cluster size distribution with the image-derived data. Due to the rarity of large clusters and to limited data, the experimental distribution is severely undersampled. It shows, however, qualitative agreement with the model prediction (Fig. S12E). Finally, to confirm that our choice of the simulation timestep *τ* did not affect our parameter estimation, we decreased *τ* by a factor of 2 from 0.001 h to 0.0005 h and found no change in the best-fit cluster size distribution within sampling uncertainties (Fig. S12F). Because our parameter grid used in the fit was coarse, we estimate the uncertainty of our best-fit parameters as the grid spacing. Our uncertainty values are therefore likely overestimated.

---

**Figure S1.**
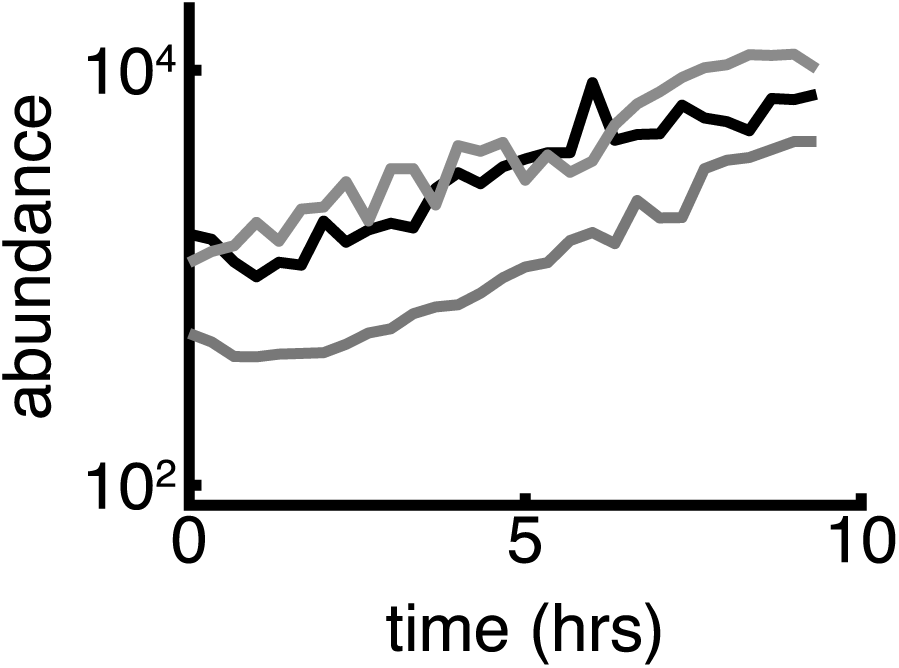
Measurement of *Enterobacter* growth rate. Image-derived quantification of initial growth dynamics in three zebrafish hosts. Imaging began approximately 8 hours after inoculation.

**Figure S2.**
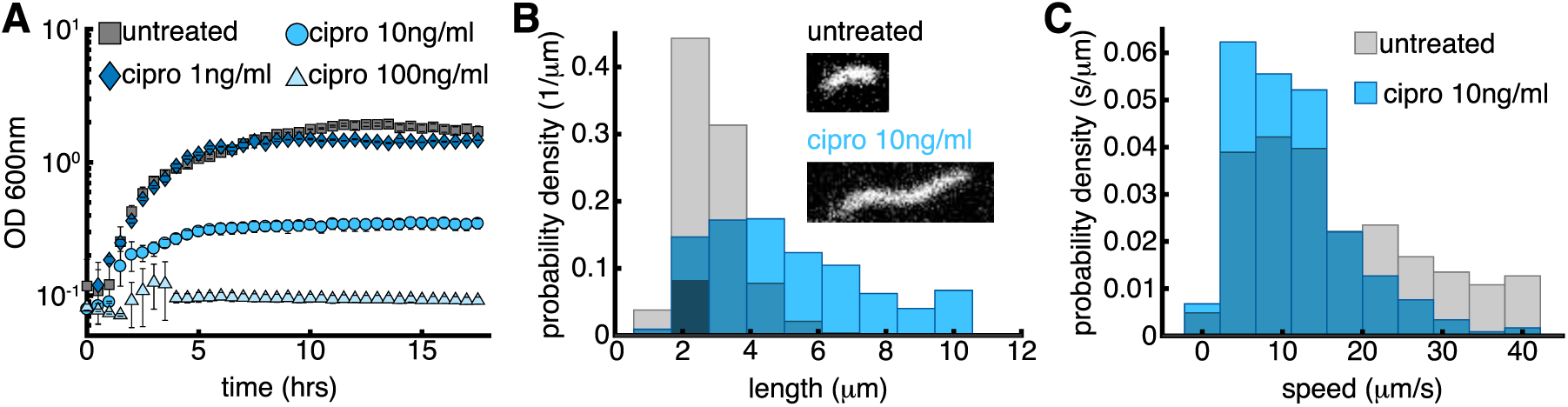
In vitro characterization of *Vibrio* response to ciprofloxacin. A: In vitro growth curves of *Vibrio* in rich media (lysogeny broth) with different ciprofloxacin concentrations. B-C: Effects of ciprofloxacin on *Vibrio* cell length and speed, with grey indicating experiments without antibiotic treatment and blue indicating exposure to 10 ng/ml ciprofloxacin. B: Distribution of *Vibrio* cell lengths. Insets show representative fluorescence microscopy images of untreated and 10 ng/ml ciprofloxacin-treated cells; inset heights = 3.5 *µ*m. C: Distribution of in vitro swimming speeds of individual bacteria.

**Figure S3.**
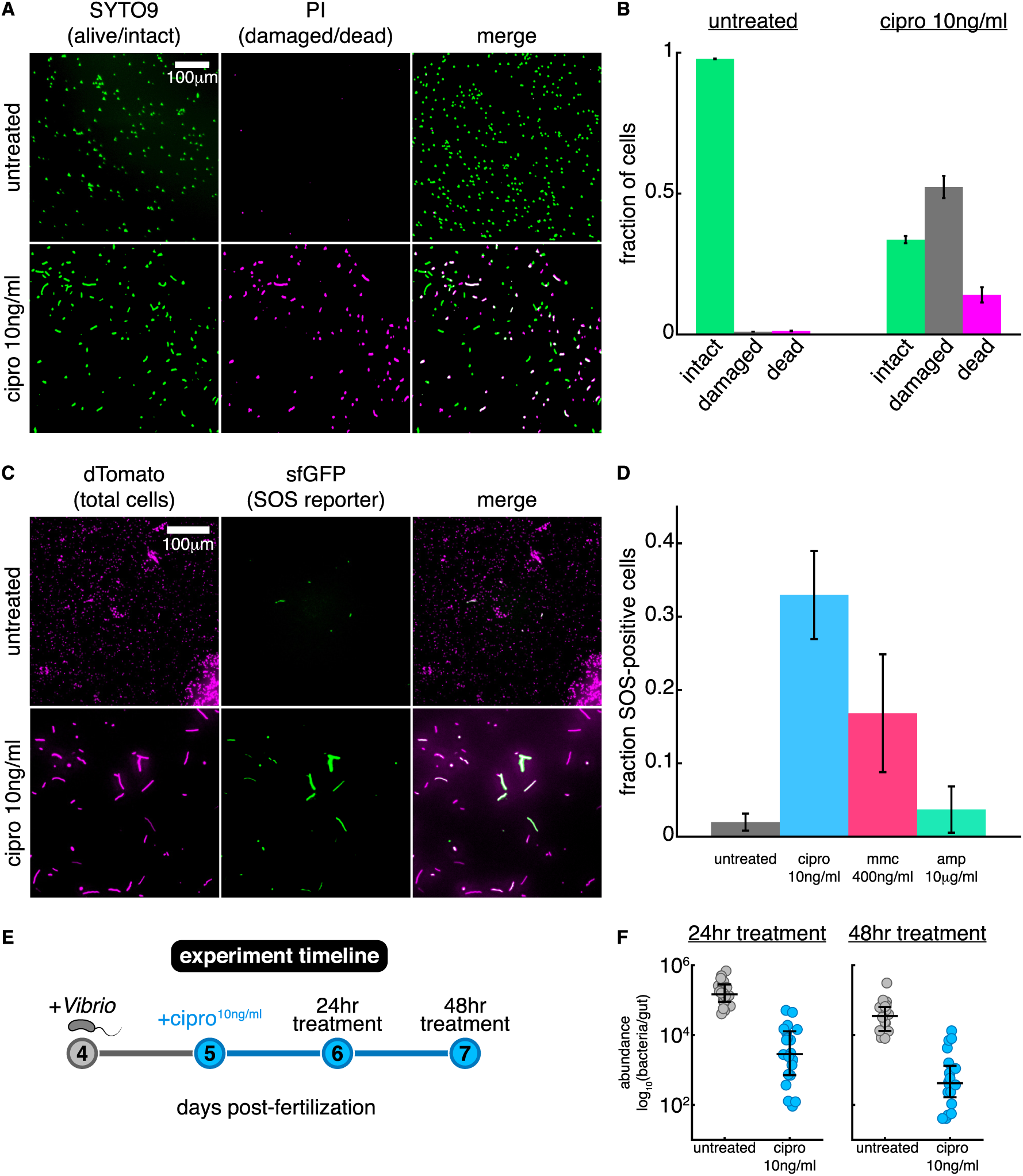
Additional *Vibrio* data. A: Representative masks of fluorescence microscopy images of in vitro viability staining. Top row, untreated, bottom row, 10 ng/ml ciprofloxacin-treated cells (6 hour treatment). SYTO9, shown in green (left panel), indicates intact cells, propridium iodine (PI), shown in magenta (middle panel), indicates dead cells. Double positive cells indicate damaged but viable cells [48], shown in white in the merged, right panel. Scale bar = 100 *µ*m. B: Quantification of in vitro viability staining by fraction of cells corresponding to each case. Mean and standard deviation across 2 replicates shown. C: Representative fluorescence microscopy images of the SOS response in untreated (top row) and 10 ng/ml ciprofloxacin treated (bottom) cells. Constitutive dTom expression is shown in magenta (left), *recN* -linked GFP expression in green (middle), merged images shown in right panel. Scale bar = 50 *µ*m. D: Quantification of SOS response in fraction of SOS+ cells (Materials and Methods), mean and standard deviations shown, *n >* 4 per treatment, total number of bacteria *>* 120 cells per treatment. E: Timeline of in vivo antibiotic treatment. F: In vivo abundances of untreated and 10 ng/ml ciprofloxacin-treated cohorts by day. Each small circle corresponds to a single host, black lines indicate medians and quartiles.

**Figure S4.**
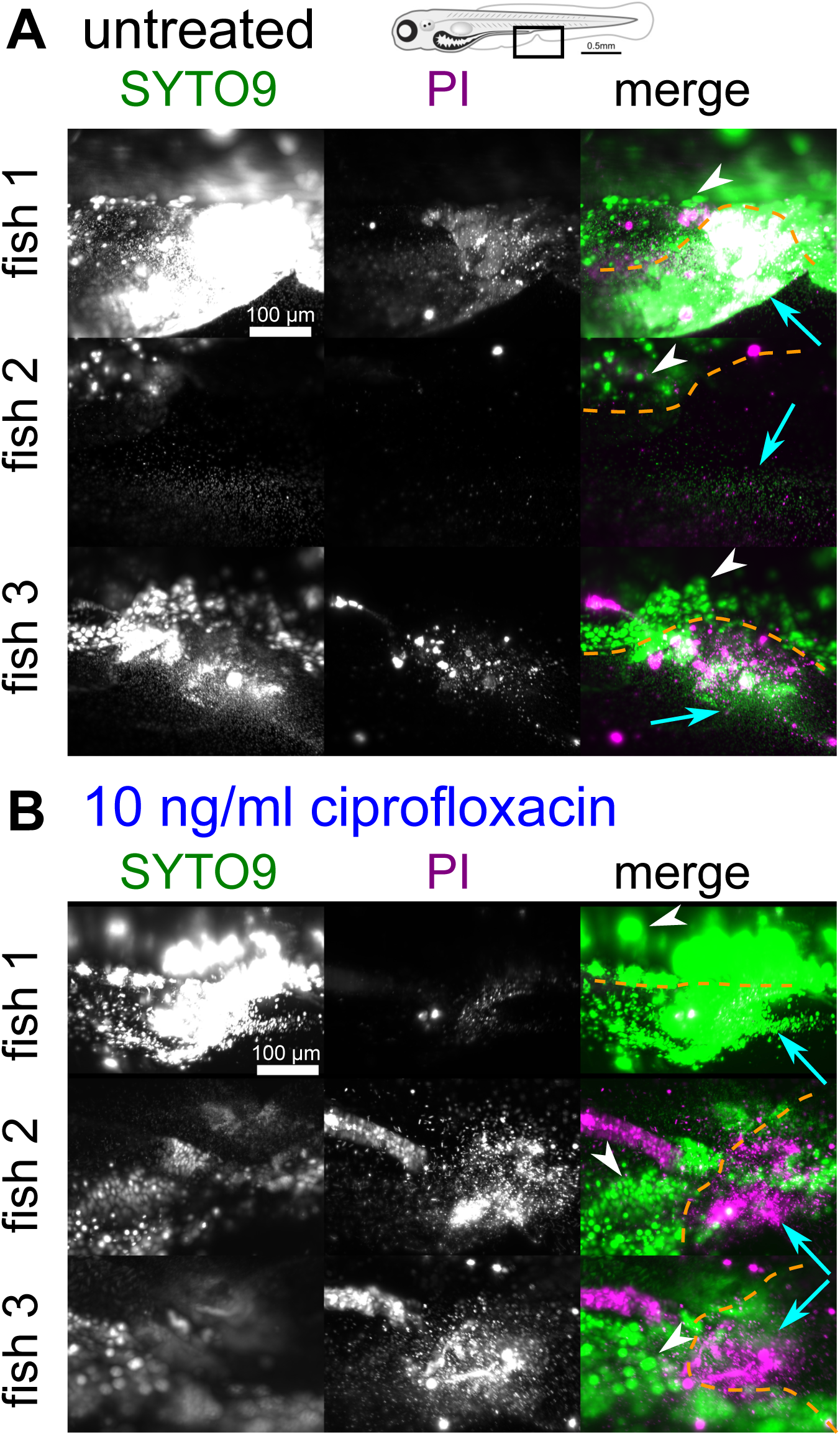
Viability staining of *Vibrio* cells expelled from the gut shows that ciprofloxacin does not induce widespread bacterial death in vivo. Three examples of fish stained with SYTO9, which indicates live bacteria, and propidium iodide (PI), which indicates dead bacteria, for both untreated (A) and 10 ng/ml ciprofloxacin-treated (B) *Vibrio*. Images were obtained by light sheet fluorescence microscopy and are maximum intensity projections of 3D images stacks. The field of view is around the vent region, as shown in the fish schematic at the top of the figure. The approximate boundary of the fish is indicated by the dashed orange line. Zebrafish cells also stain and constitute the bulk of the fluorescence in the images. Examples of zebrafish cells are indicated by white arrow heads. Examples of bacterial cells are indicated by the cyan arrows.

**Figure S5.**
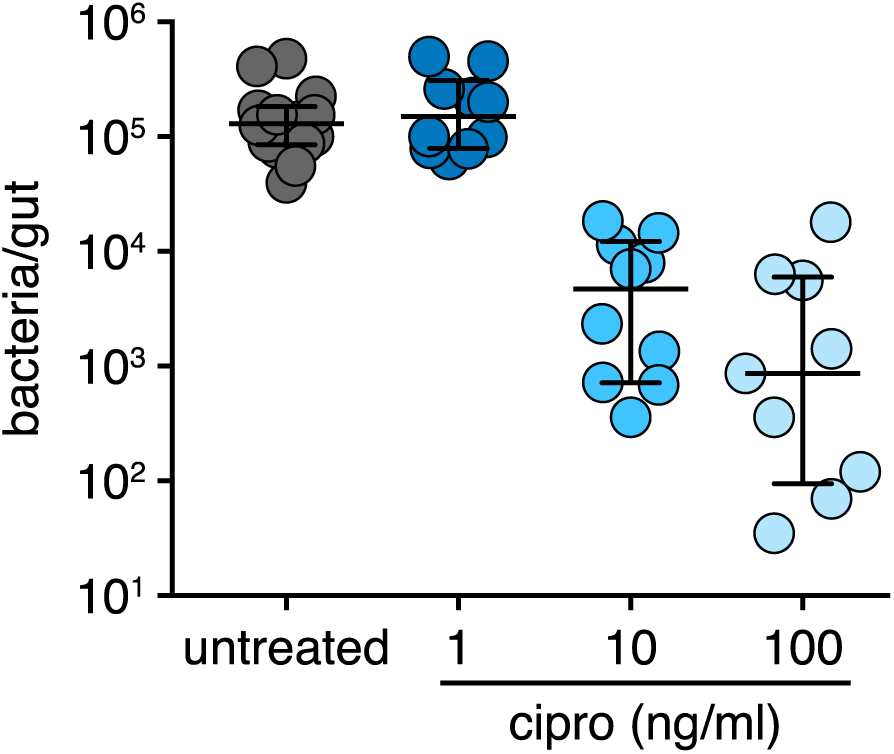
In vivo ciprofloxacin dose response for *Vibrio*. *Vibrio* was mono-associated with germfree larval zebrafish for 24 hours prior to being left untreated, or treated with either 1, 10, or 100 ng/ml ciprofloxacin for an additional 24 hours. *Vibrio* abundances were determined by dissection and plating. Each circle corresponds to a single host intestine, black lines indicate medians and quartiles. Data for the ‘untreated’ and ‘cipro 10 ng/ml’ groups were included in Figure 2D and Supplemental Figure 2F, where they were combined with repeated experiments.

**Figure S6.**
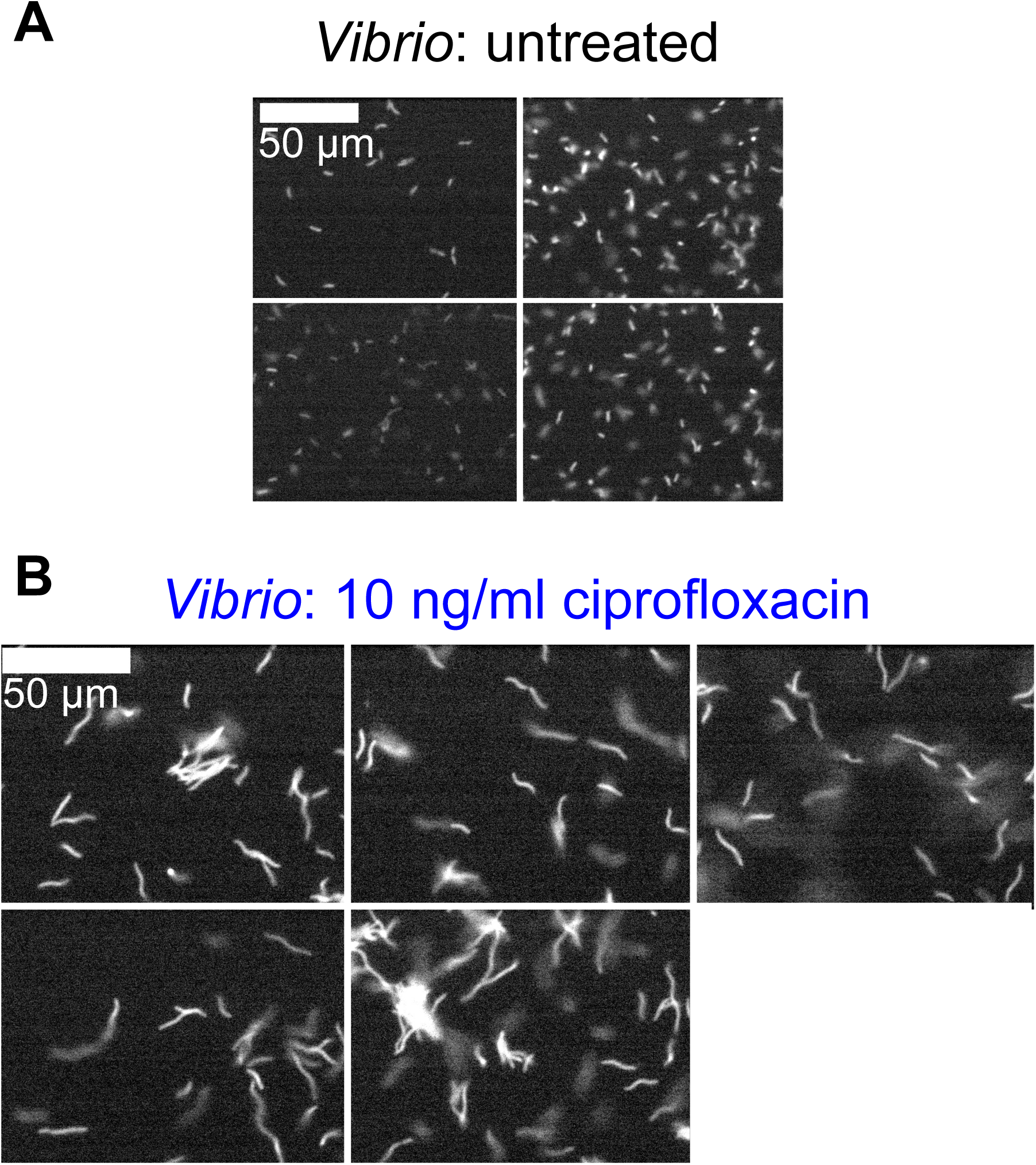
*Vibrio* does not form large aggregates in vitro in response to ciprofloxacin. Representative fluorescence microscopy images of untreated (A) and 10 ng/ml ciprofloxacin-treated (B) *Vibrio* cells. Sample preparation and treatment are described in the *Cell length and swimming speed* portion of the Materials and Methods section.

**Figure S7.**
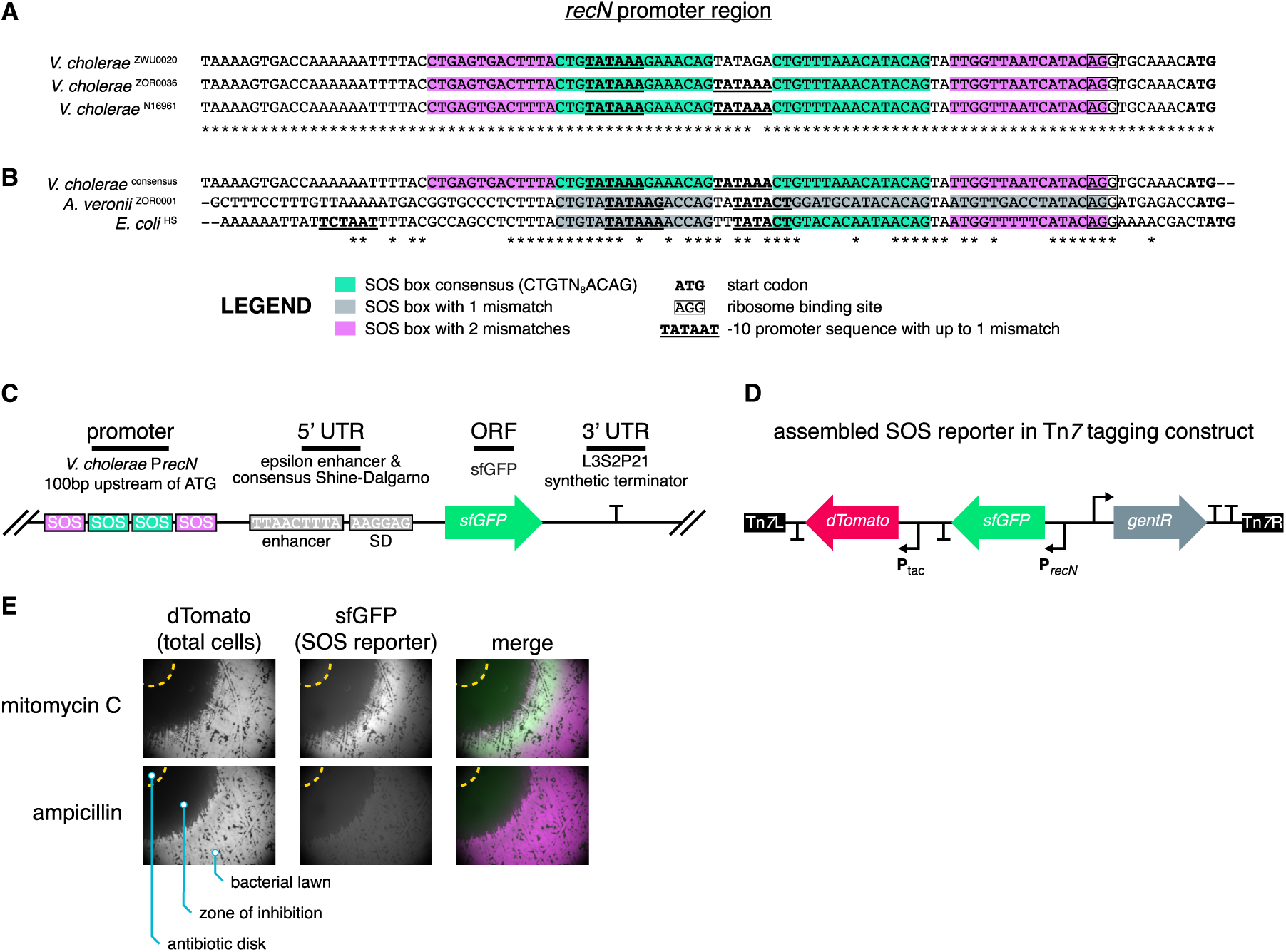
Design and construction of fluorescent SOS reporter. A: Alignment of 100bp *recN*promoter region plus start codon for the closely related *V. cholerae* strains ZWU0020 (zebrafish isolate used in this study, IMG gene ID: 2705597027, locus tag: ZWU0020 01601), ZOR0036 (zebrafish isolate, IMG gene ID: 2705599600, locus tag: ZOR0036 00266), and El Tor N16961 (human pandemic isolate, IMG gene ID: 637047325, locus tag: VC0852). B: Alignment of 100bp *recN* promoter region plus start codon of the *V. cholerae* consensus *recN* promoter, *Aeromonas veronii* (zebrafish isolate, IMG gene ID: 2526373590, locus tag: L972 03073), and *E. coli* HS (human commensal isolate IMG gene ID: 640921890, locus tag: EcHS A2774). For panels A and B, SOS boxes are shaded based on their conservation to the consensus gammaproteobacterial sequence (CTGTN_8_ACAG); ‘ATG’ start codons are bolded; putative ribosome binding sites are boxed; and putative, near-consensus -10 promoter sequences (TATAAT) are bolded and underlined. C: Schematic of *recN* -based fluorescent SOS reporter. Promoter comprises the consensus *V. cholerae recN* promoter region (P*recN*), which was derived from the sequence alignment in panel A. The synthetic 5’ untranslated region (UTR) contains an epsilon enhancer and consensus Shine-Dalgarno sequence. The open reading frame (ORF) encodes superfolder green fluorescent protein (sfGFP). And the 3’ UTR contains the synthetic transcriptional terminator L3S2P21. D: Schematic of assembled SOS reporter in the context of the Tn*7* tagging construct. Tn*7* L and Tn*7* R inverted repeats flank the Tn*7* transposon. The SOS reporter was inserted upstream of a dTomato gene that is constitutively expressed from a synthetic Ptac promoter. A gene encoding gentamicin resistance (*gentR*) was used to facilitate genetic manipulation. E: Disk diffusion assays verifying SOS reporter activity. *Vibrio* ZWU0020 carrying the SOS reporter was spread onto agar plates using glass beads at a density that would give rise to a lawn of growth. Circular disks of Whatman filter paper (amber dashed lines) loaded with either the genotoxic agent mitomycin C or the cell wall-targeting beta-lactam antibiotic ampicillin were then placed in the center of the agar plates. After overnight incubation at 30*^◦^*C, plates were imaged using a fluorescence stereomicroscope. In the presence of mitomycin C, cells adjacent to the zone of inhibition (i.e., the area where there is no bacterial growth) robustly expressed sfGFP whereas in the presence of ampicillin they did not.

**Figure S8.**
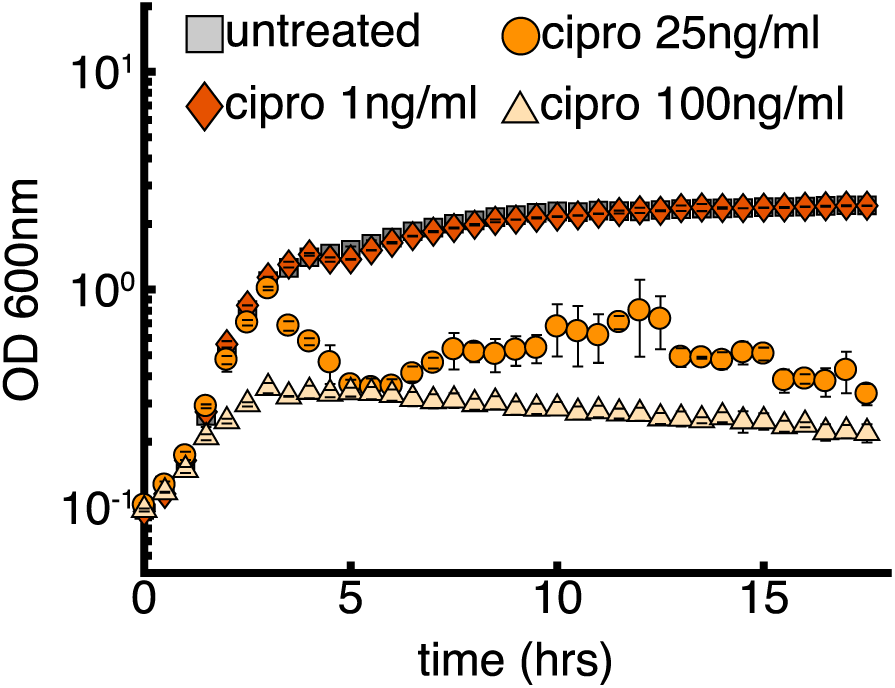
In vitro growth curves (in lysogeny broth) of *Enterobacter* with varying concentrations of ciprofloxacin.

**Figure S9.**
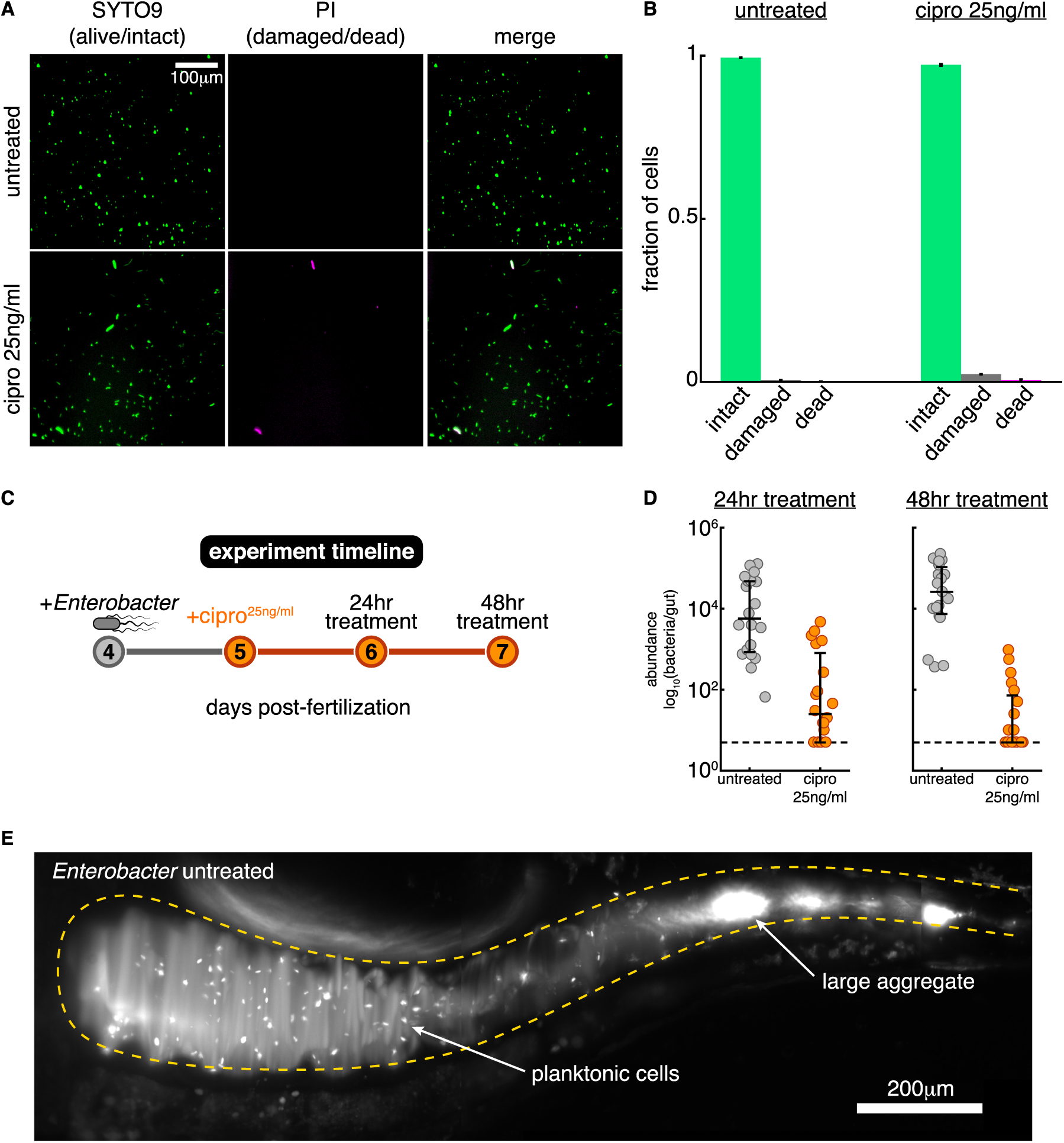
Additional *Enterobacter* data. A: Representative fluorescence microscopy images of in vitro viability staining. Top row, untreated, bottom row, 25 ng/ml ciprofloxacin-treated cells (6 hour treatment). SYTO9, shown in green (left panel), indicates intact cells, propridium iodine (PI), shown in magenta (middle panel), indicates dead cells. Double positive cells indicate damaged but viable cells [48], shown in white in the merged, right panel. Scale bar = 100 *µ*m. B: Quantification of in vitro viability staining by fraction of cells corresponding to each case. Mean and standard deviation across 2 replicates shown. C: Timeline of in vivo antibiotic treatment. D: In vivo abundances of untreated and 25 ng/ml ciprofloxacin-treated cohorts by day. Each small circle corresponds to a single host, black lines indicate medians and quartiles. E: Maximum intensity projection of untreated *Enterobacter* population showing an example of a population containing a single large cluster and several single cells. Scale bar = 200 *µ*m.

**Figure S10.**
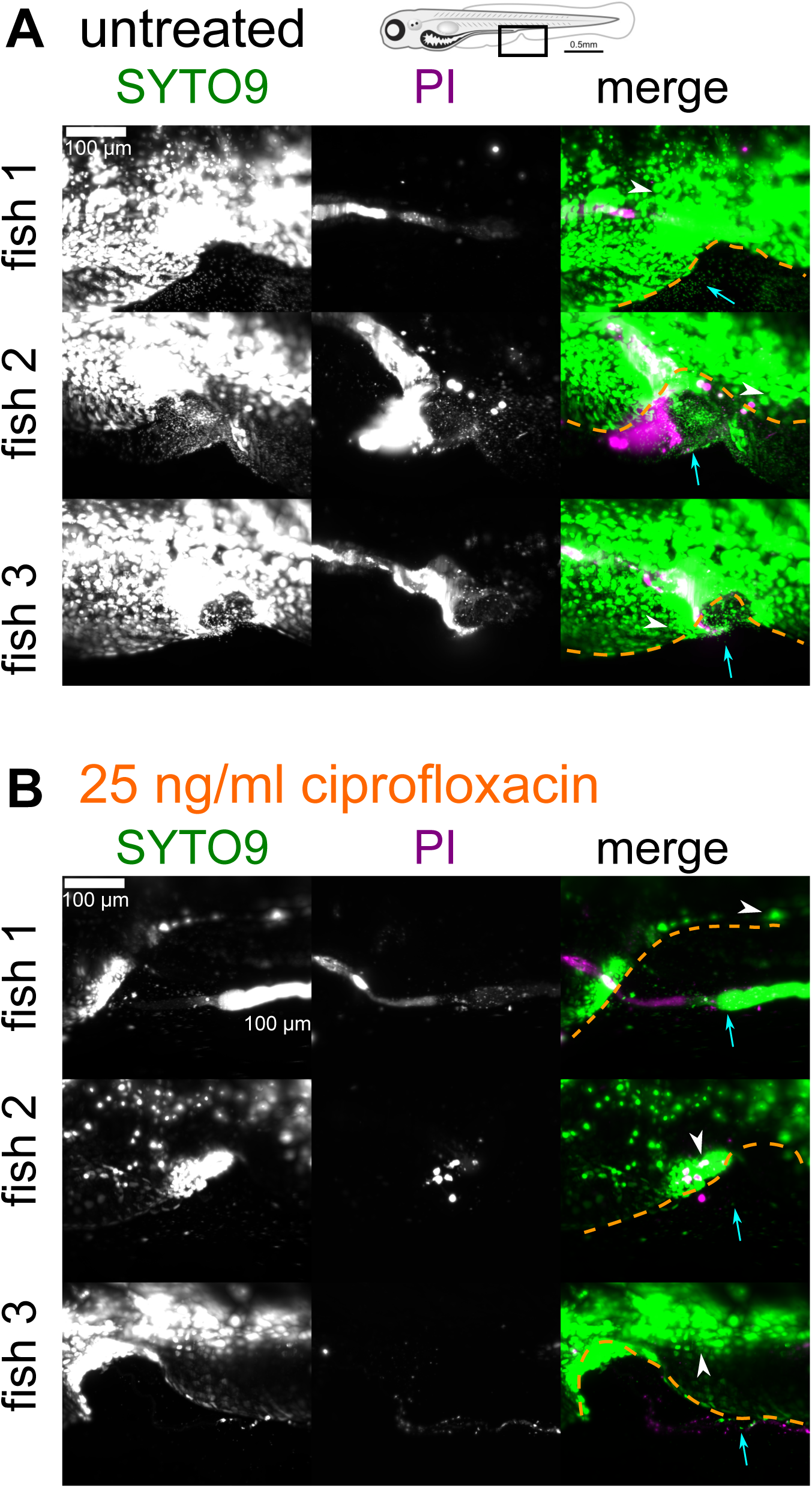
Viability staining of *Enterobacter* cells expelled from the gut shows that ciprofloxacin does not induce widespread bacterial death in vivo. Three examples of fish stained with SYTO9, which indicates live bacteria, and propidium iodide (PI), which indicates dead bacteria, for both untreated (A) and 25 ng/ml ciprofloxacin-treated (B) *Enterobacter*. Images were obtained by light sheet fluorescence microscopy and are maximum intensity projections of 3D images stacks. The field of view is around the vent region, as shown in the fish schematic at the top of the figure. The approximate boundary of the fish is indicated by the dashed orange line. Zebrafish cells also stain and constitute the bulk of the fluorescence in the images. Examples of zebrafish cells are indicated by white arrow heads. Examples of bacterial cells are indicated by the cyan arrows.

**Figure S11.**
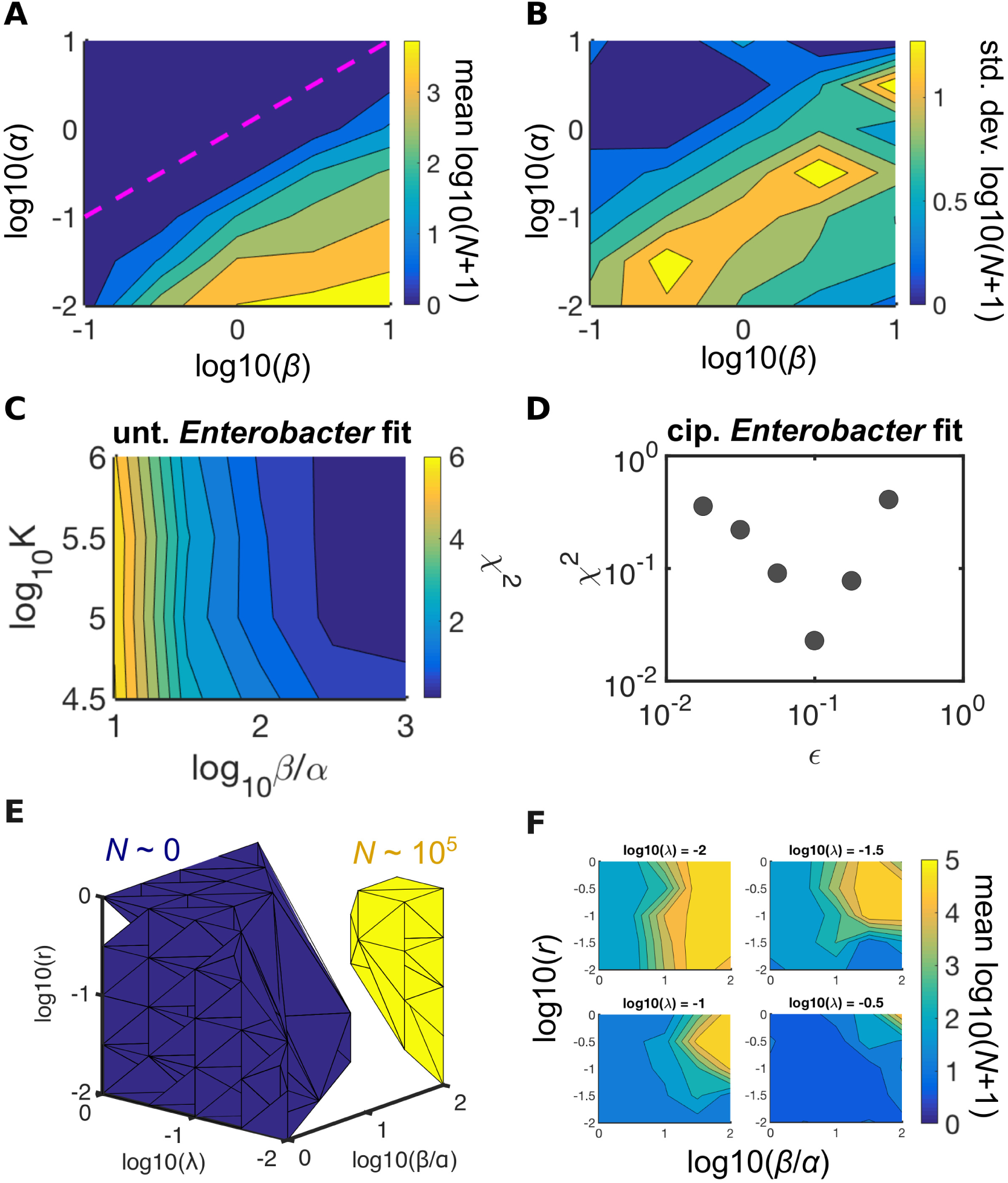
Additional model details. A-B: Simulated heatmap of mean (A) and standard deviation (B) of log_10_(abundance + 1) for varying values of aggregation and fragmentation rates. Both mean and standard deviation depend primarily on the ratio of fragmentation to aggregation rates, rather than on each rate independently. Dashed magenta line in (A) represents *α* = *β*. Parameters: *r* = 0.27 hr^−1^, *λ* = 0.11 hr^−1^, *K* = 10^5^, simulation time = 64 hours, number of trials decreased logarithmically with *β* from 1000 to 10. Units of *α* and *β* are hr^−1^. C: Heatmap of *χ*^2^ for untreated *Enterobacter* fit to 7 dpf abundances (Materials and Methods). Parameters: *r* = 0.27 hr^−1^, *λ* = 0.11 hr^−1^, *α* = 0.1 hr^−1^, simulation time = 64 hours, number of trials decreased logarithmically with *β* from 1000 to 10. D: *χ*^2^ for fit to 6 dpf ciprofloxacin-treated *Enterobacter* abundances as a function of the scaling parameter *∊*, which scales the growth and fragmentation rates simultaneously according to *r → ∊r* and *β → ∊β*. A clear minimum is seen at *∊* = 0.1. Parameters: *r* = 0.27 hr^−1^, *λ* = 0.11 hr^−1^, *α* = 0.1 hr^−1^, *β* = 10^1.5^ hr^−1^, simulation time = 64 hours, number of trials decreased logarithmically with *β* from 1000 to 10. E: 3D phase diagram of log_10_(abundance+ 1) with axes fragmentation/aggregation (*β/α*), growth rate (*r*), and expulsion rate (*λ*). Blue isosurface represents log_10_(abundance + 1) = 0.5 *±* 0.5, yellow isosurface represents log_10_(abundance + 1) = 5.5 *±* 0.5. Parameters: *α* = 0.1 hr^−1^, simulation time = 64 hours, number of trials decreased logarithmically with *β* from 1000 to 10. Units of *α* and *β* are hr^−1^. F: Slices through the 3D phase diagram in (E) for different values of *λ*.

**Figure S12.**
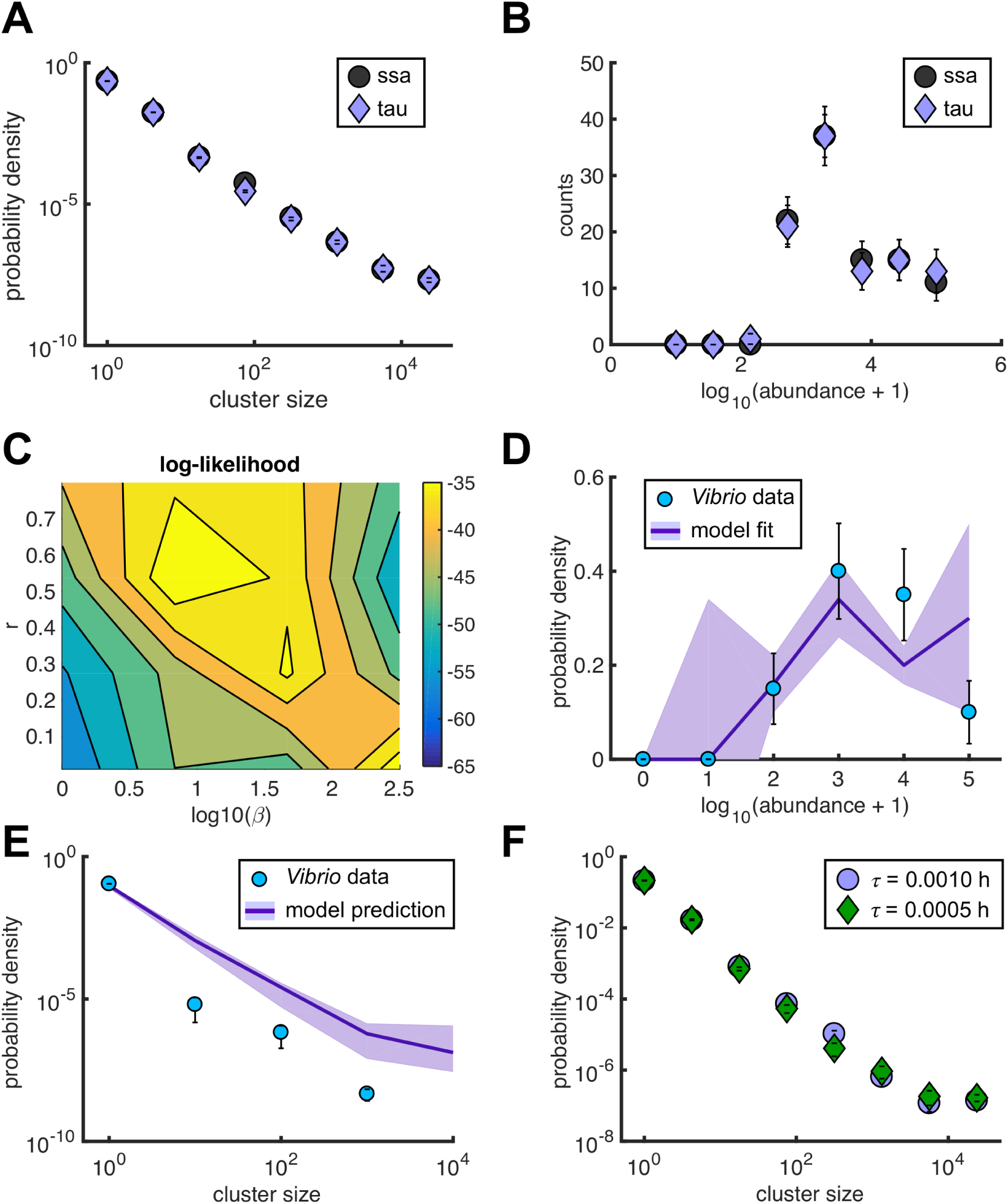
Tau leaping simulations and *Vibrio* parameter inference. A-B: Comparison of direct stochastic simulation (“ssa”, gray circles) and our fixed-tau leaping (“tau”, purple diamonds) algorithm with *τ* = 0.001 h. Simulations using both methods were run with the best-fit parameters for untreated *Enterobacter* and 100 replicates. Both the cluster size distribution (A) and abundance histogram (B) show excellent agreement between the two methods. C-E: Details of model fit to ciprofloxacin-treated *Vibrio* 24 h abundances. C: Heat map of log-likelihood. A manual grid search was performed over growth rate (*r*) and fragmentation rate (*β*). D: Comparison of the best-fit abundance distribution (purple line) to experimental data (blue circles). E: Comparison of the predicted cluster size distribution (purple line) to experimental data (blue circles). Here, all model parameters were fixed at their previously determined, best-fit values; there were no additional free parameters. The experimental data distribution is severely undersampled, estimated from just 4 fish. F: Confirmation that the best-fit solution is independent of our choice of *τ*, indicating that simulations were performed with sufficient resolution. Simulations were run with the best-fit parameters but with *τ* decreased by a factor of 2, from *τ* = 0.001 h (purple circles) to *τ* = 0.0005 h (green diamonds). Distributions agree with one another within sampling uncertainties.

**Figure S13.**
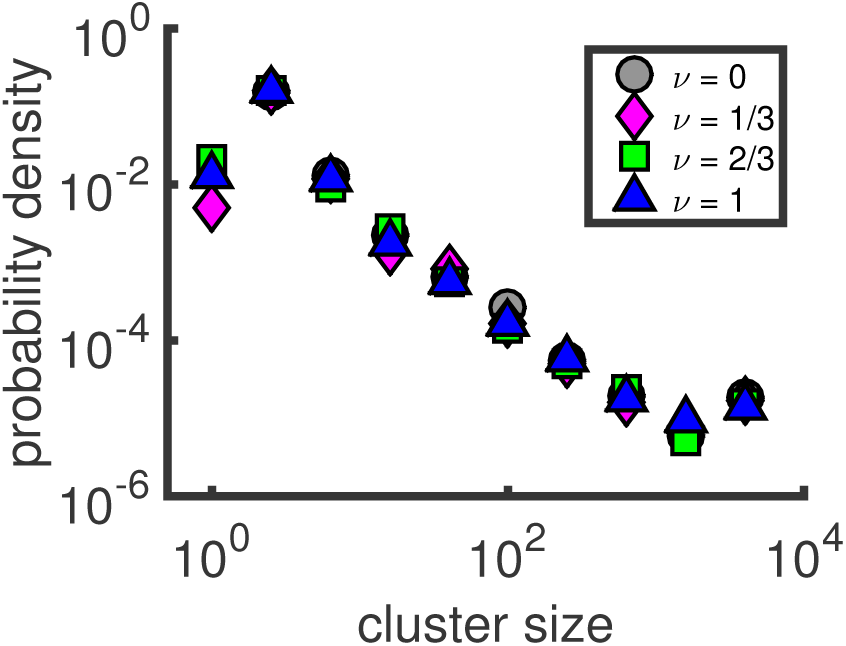
Model cluster size distributions are independent of how expulsion rate scales with cluster size. Simulations were run with the expulsion rate depending on cluster size according to *E_n_* = *λ*(*n/K*)*^ν^*, with *K* the carrying capacity, and the exponent *ν* was varied. This ansatz keeps the expulsion rate of clusters of size *K* constant. The resulting cluster size distributions agree with one another within sampling uncertainties, which are smaller than the marker size. This result justifies our use of the simple constant form of the expulsion kernel, *E_n_* = *λ*.

## Supplemental Movie captions

**Supplemental Movie 1**

Light sheet fluorescence microscopy movie of untreated *Vibrio* swimming in a 6 dpf zebrafish gut. The density of cells is highest on the left (anterior), where single cells cannot be resolved and the population appears as a single bright region (see also Figure 1C). On the right (posterior), single cells are more easily resolved and are seen swimming in and out of the intestinal folds. Each frame is from the same optical plane. Scale bar = 50 *µ*m.

**Supplemental Movie 2**

Animated z-stack of light sheet fluorescence microscopy images of untreated *Enterobacter* in a 6 dpf zebrafish gut. Bacterial clusters (bright white puncta) of diverse sizes are evident, from single cells up to a single cluster containing thousands of cells that appears at a z depth of *∼* 70 *µ*m. Hazy reflection of light off of the fish’s swim bladder can be seen outside the intestinal boundary in the upper right section of the images. Scale bar = 50 *µ*m.

**Supplemental Movie 3**

Fluorescence microscopy movie of untreated *Vibrio* swimming between a glass slide and a coverslip (Materials and Methods). Scale bar = 20 *µ*m.

**Supplemental Movie 4**

Fluorescence microscopy movie of *Vibrio* treated with 10 ng/ml ciprofloxacin swimming between a glass slide and a coverslip (Materials and Methods). Cells have undergone filamentation. Scale bar = 20 *µ*m.

**Supplemental Movie 5**

Time-lapse light sheet fluorescence microscopy movie of an established *Vibrio* population responding to 10 ng/ml ciprofloxacin. Each frame is a maximum intensity projection of the full 3D intestinal volume. The time between frames is 20 min. Initially, the population consists of a dense collection of individual, motile cells (Supplemental Movie 1, Figure 1C). Antibiotics are added after the second frame of the movie. Following motility loss, cells leave the swarm and are compacted into aggregates, which are subject to strong transport down the length of the intestine and are eventually expelled. Scale bar = 200 *µ*m.

**Supplemental Movie 6**

Light sheet fluorescence microscopy movies of *Vibrio* in fish treated with 10 ng/ml ciprofloxacin. The left panel movie shows constitutive dTom expression. The right panel movie was taken immediately after the left panel movie and shows a GFP reporter of the SOS response (Materials and Methods), which is expressed in cells strongly affected by ciprofloxacin (Fig. S3C and S3D). GFP-positive cells swim slowly or are aggregated. Each frame is from the same optical plane. Scale bar = 25 *µ*m.

**Supplemental Movie 7**

Time-lapse light sheet fluorescence microscopy movie of an untreated *Enterobacter* population showing an example of the expulsion process. Each frame is a maximum intensity projection of the full 3D intestinal volume. Time between frames is 10 min. The population is initially comprised of many small bacterial clusters and a single large cluster. Over time, small clusters are incorporated into the large one and the mass is transported down the length of the gut and expelled. Image intensities are log-transformed. Scale bar = 200 *µ*m.

**Supplemental Movie 8**

Time-lapse light sheet fluorescence microscopy movie of an untreated *Enterobacter* population showing an example of the aggregation process. Each frame is a maximum intensity projection of the full 3D intestinal volume. Time between frames is 10 min. A collection of initially disconnected bacterial clusters on the left (anterior) side of the field of view gradually combine into a single cluster. Image intensities are log-transformed. Scale bar = 200 *µ*m.

**Supplemental Movie 9**

Time-lapse light sheet fluorescence microscopy movie of an untreated *Enterobacter* population showing examples of the growth and fragmentation processes. Each frame is a maximum intensity projection of the full 3D intestinal volume. The time between frames is 20 min. The movie begins 8 hours after the initial exposure to *Enterobacter*, by which time a small founding population has been established. Over time, the aggregates grow in size as cells divide, and new single cells also appear in the vicinity of the aggregates, likely due to fragmentation. Individual cell divisions from planktonic cells are also visible. Image intensities are log-transformed. Scale bar = 200 *µ*m.

**Supplemental Movie 10**

Light sheet fluorescence microscopy movie of *Vibrio* in a fish treated with 10 ng/ml ciprofloxacin for *∼*18 hours. Each frame is from the same optical plane, which spans the anterior-most region of the intestine known as the intestinal bulb (Fig. 1B). The bright signal in the left (anterior) side of the frame is a dense, motile swarm of planktonic cells (Supplemental Movie 1 and Fig. 1C). Moving from left to right (anterior-posterior) across the field of view, cells exhibiting filamentation and reduced motility are evident, along with the beginnings of small aggregates. Scale bar = 50 *µ*m.

**Supplemental Movie 11**

Light sheet fluorescence microsocopy movie of *Vibrio* in a fish treated with 10 ng/ml ciprofloxacin for *∼*18 hours. Each frame from the same single optical plane that captures a portion of the midgut (Fig. 1B). The bright signal is an aggregate of *Vibrio* cells that nearly fills the width of the midgut lumen. Two cells are seen swimming near the end of the movie. Scale bar = 25 *µ*m.

