## Supplemental Data Files for "Sublethal antibiotics collapse gut bacterial populations by enhancing aggregation and expulsion": README.txt

README.txt

Information for Supplemental Data Files for paper, “Sublethal antibiotics collapse gut bacterial populations by enhancing aggregation and expulsion”. For more information, please contact Brandon Schlomann at, or Raghuveer Parthasarathy at. 

—BS

%%%%%In vitro data%%%%%%%%%%
abx_growth_curve_data.xlsx - OD600 measurements that generated Figures S2A and S8.

vibrio_length_data.csv - lengths of Vibrio cells, top row = untreated, bottom row = cip10

vibrio_speed_data.csv - speeds of Vibrio cells, top row = untreated, bottom row = cip10

abx_invitro_livedead_data.xlsx - fractions of intact/damaged/dead cells from in vitro live/dead staining in Fig S3B. Details in file.

vibrio_sos_invitro_abx.xlsx - fractions of sos positive Vibrio cells from in vitro characterization. Details in file.

%%%%%%Plating data%%%%%%%%%%
abx_plating_data.xlsx - CFUs from in vivo abundance measurements that generated Figures 2C,D 3A,B, and S5.

abx_water_plating_data.xlsx - CFUs of flask water abundances from Figures 2C and 3A.

normalized_cfus.xlsx - normalized abundances plotted in figs 2C and 3A.

%%%%%%In vivo time series data%%%%%%%%%
ent_initial_growth_timeseries.csv (Fig 4D), ent_main_abundance_timeseries.csv (Fig 4C), vibrio_untreated_abundance_timeseries.csv (Fig 2H), vibrio_cip_abundance_timeseries.csv (Fig 2H), vibrio_untreated_com_timeseries.csv (Fig 2G), vibrio_cip_com_timeseries.csv (Fig 2G)

These files contain the data for imaging-derived measurements of abundance and center of mass in untreated and treated cases, and all have the same structure. Each row corresponds to a host and contains a comma separated list of first, the time points, and second, the measurements. 

%%%%%%%Vibrio in vivo 1D distirbutions (Fig 2E)%%%%%%%%%%%%
vibrio_unt_binned_1D_dists.csv - untreated 1D distributions

vibrio_cip_binned_1D_dists.csv - cip10 1D distributions

%%%%%%%%%%In vivo Enterobacter cluster sizes (Figs 3C, 5A)%%%%%%%%
enterobacter_untreated_clusters.csv - each row corresponds to an untreated host and contains a list of bacterial cluster sizes

enterobacter_cip_clusters.csv - each row corresponds to a cip25 treated host and contains a list of  bacterial cluster sizes
